# The phylogenetic range of bacterial and viral pathogens of vertebrates

**DOI:** 10.1101/670315

**Authors:** Liam P. Shaw, Alethea D. Wang, David Dylus, Magda Meier, Grega Pogacnik, Christophe Dessimoz, Francois Balloux

## Abstract

Many major human pathogens are multi-host pathogens, able to infect other vertebrate species. Describing the general patterns of host-pathogen associations across pathogen taxa is therefore important to understand risk factors for human disease emergence. However, there is a lack of comprehensive curated databases for this purpose, with most previous efforts focusing on viruses. Here, we report the largest manually compiled host-pathogen association database, covering 2,595 bacteria and viruses infecting 2,656 vertebrate hosts. We also build a tree for host species using nine mitochondrial genes, giving a quantitative measure of the phylogenetic similarity of hosts. We find that the majority of bacteria and viruses are specialists infecting only a single host species, with bacteria having a significantly higher proportion of specialists compared to viruses. Conversely, multi-host viruses have a more restricted host range than multi-host bacteria. We perform multiple analyses of factors associated with pathogen richness per host species and the pathogen traits associated with greater host range and zoonotic potential. We show that factors previously identified as important for zoonotic potential in viruses—such as phylogenetic range, research effort, and being vector-borne—are also predictive in bacteria. We find that the fraction of pathogens shared between two hosts decreases with the phylogenetic distance between them. Our results suggest that host phylogenetic similarity is the primary factor for host-switching in pathogens.

## Introduction

Pathogens vary considerably in their host ranges. Some can only infect a single host species, whereas others are capable of infecting a multitude of different hosts distributed across diverse taxonomic groups. Multi-host pathogens have been responsible for the majority of recent emerging infectious diseases in both human (Jones et al., 2008; Karesh et al., 2012; Taylor, Latham, & Woolhouse, 2001; Woolhouse & Gowtage-Sequeria, 2005) and animal populations (Cleaveland, Laurenson, & Taylor, 2001; Daszak, Cunningham, & Hyatt, 2000). A number of studies have concluded that pathogens having a broad host range spanning several taxonomic host orders constitute a higher risk of disease emergence, compared to pathogens with more restricted host ranges (Cleaveland et al., 2001; Howard & Fletcher, 2012; Kreuder Johnson et al., 2015; McIntyre et al., 2014; Olival et al., 2017; Parrish et al., 2008; Woolhouse & Gowtage-Sequeria, 2005).

An important biological factor that is likely to limit pathogen host-switching is the degree of phylogenetic relatedness between the original and new host species. For a pathogen, closely related host species can be considered akin to similar environments, sharing conserved immune mechanisms or cell receptors, which increases the likelihood of pathogen ‘pre-adaptation’ to a novel host. Barriers to infection will depend on the physiological similarity between original and potential host species (Poulin & Mouillot, 2005), factors that can depend strongly on host phylogeny. Indeed, the idea that pathogens are more likely to switch between closely related host species has been supported by studies in several host-pathogen systems (Davies & Pedersen, 2008; Faria, Suchard, Rambaut, Streicker, & Lemey, 2013; Streicker et al., 2010; Waxman, Weinert, & Welch, 2014). The likelihood of infection of a target host has also been found to increase as a function of phylogenetic distance from the original host in a number of experimental infection studies (Gilbert & Webb, 2007; Longdon, Hadfield, Webster, Obbard, & Jiggins, 2011; Perlman & Jaenike, 2003; Russell et al., 2009).

Nevertheless, there are also numerous cases of pathogens switching host over great phylogenetic distances, including within the host-pathogen systems mentioned above. For example, a number of generalist primate pathogens are also capable of infecting more distantly-related primates than expected (Cooper et al., 2012). Moreover, for zoonotic diseases, a significant fraction of pathogens have host ranges that encompass several mammalian orders, and even non-mammals (Woolhouse & Gowtage-Sequeria, 2005). Interestingly, host jumps over greater phylogenetic distances may lead to more severe disease and higher mortality (Farrell & Davies, 2019). One factor that could explain why transmission into more distantly related new hosts occurs at all is infection susceptibility; some host clades may simply be more generally susceptible to pathogens (e.g. if they lack broad resistance mechanisms). Pathogens would therefore be able to jump more frequently into new hosts in these clades regardless of their phylogenetic distance from the original host. In support of this, experimental cross-infections have demonstrated that sigma virus infection success varies between different *Drosophila* clades (Longdon et al., 2011), and a survey of viral pathogens and their mammalian hosts found that host order was a significant predictor of disease status (Levinson et al., 2013).

While an increasing number of studies have described broad patterns of host range for various pathogens (see Table 1), most report only crude estimates of the breadth of host range, and there have been few attempts to systematically gather quantitatively explicit data on pathogen host ranges. As noted by Bonneaud, Weinert, & Kuijper (2019), most work on pathogen emergence has focused on viruses since these are the cause of many high-profile outbreaks (e.g. *Ebolavirus*), but it is plausible that the processes underlying emergence may be different in bacterial pathogens. We thus have little understanding of the overall variation in host range both within and amongst groups of pathogens. This has limited our ability to examine how pathogen host range correlates with the emergence of infectious diseases. Here, we address this gap in the literature by considering both bacterial and viral pathogens in the same dataset.

**Table 1.** Notable previous studies of the host range of pathogens using a database approach.

| Study | Database details | Methods | Pathogen types | Host range classification | Main conclusions |
| --- | --- | --- | --- | --- | --- |
| **Taylor et al. (2001)** | 1,415 human pathogens | Literature review | Viruses and prions, bacteria and rickettsia, fungi, protozoa, helminths | Categorical: zoonotic or not | Found that 61% of human pathogens are zoonotic.<br>First study identifying zoonotic pathogens as a major risk factor for human disease emergence. |
| **Cleaveland et al. (2001)** | 1,922 human and domestic mammal (livestock and carnivore) pathogens | Literature review | Viruses and prions, bacteria and rickettsia, fungi, protozoa, helminths | Categorical: taxonomic grouping of mammal hosts (carnivores, ungulates, primates, bats, rodents and marine mammals)<br>Simple quantitative: single or multiple host; human, domestic or wildlife hosts; hosts of one or more taxonomic orders | Found that 63% of pathogens infect multiple hosts, with multiple-host infections making up a higher proportion of domestic mammal pathogens than human pathogens.<br>First study providing simple quantitative data on the host ranges of human and domestic mammal pathogens; and identifying the ability to infect multiple hosts (especially across taxonomic orders) as a risk factor for human and domestic mammal diseases emergence. |
| **Woolhouse & Gowtage-Sequeria (2005)** | 1,407 human pathogens | Literature review | Viruses and prions, bacteria and rickettsia, fungi, protozoa, helminths | Categorical: zoonotic or not; type of nonhuman vertebrate host (broad categories: bats, carnivores, primates, rodents, ungulates, other mammals and nonmammals)<br>Simple quantitative: number of host types (0 – human only, 1, 2 or 3+ host types). | Zoonotic pathogens identified as major risk factor for human diseases emergence, with the fraction of emerging pathogen species increasing with the breadth of host range (number of host types). |
| **McIntyre et al. (2014)** | 2,597 pathogen species across 47 mammalian and avian hosts (including humans and animals commonly used in Europe as food or kept as pets)<br>4,223 host-pathogen associations | Automated data mining of NCBI meta-data and semi-automated literature searches | Viruses and prions, bacteria and rickettsia, fungi, protozoa, helminths | Categorical: taxonomic grouping of mammal hosts (carnivores, ungulates, primates, bats, rodents and marine mammals)<br>Simple quantitative: 1, 2, or 2+ host species | Pathogens having greater numbers of host species have increased odds of being a risk factor for disease emergence.<br>Multiple-host infections make up a higher proportion of domestic mammal pathogens than human pathogens. |
| **Kreuder Johnson et al. (2015)** | 162 zoonotic pathogens | Literature review | Viruses | Simple quantitative: viral host range (host plasticity) calculated as the total count of animal taxonomic orders and ecological groups recognized as hosts | Viruses with high host plasticity are more likely to amplify viral spillover by human-to-human transmission and have broader geographic spread. |
| **Wardeh et al. (2015)** | 8,905 'cargo' (pathogen) species across 6,314 'carrier' (host) species (22,515 carrier/cargo interactions) | Automated data mining of NCBI databases (Taxonomy, Nucleotide, PubMed) | Viruses and prions, bacteria and rickettsia, fungi, protozoa, helminths | Not applicable | Verified individual species interactions from automated procedures against interactions identified by literature reviews. |
| **Han et al. (2016)** | Zoonotic pathogens of 27 orders of terrestrial mammals | Literature review | Viruses, bacteria, protozoa, helminths | Categorical: mammalian host order | Identified the proportion of zoonotic host species in each mammalian order (carnivores and rodents harbor the most zoonoses).<br>Mammals carry more bacteria than any other pathogen type, followed by viruses. |
| **Olival et al. (2017)** | 586 pathogens across 754 mammal species (2,805 host-pathogen associations) | Literature and database review | Viruses | Quantitative: phylogenetic host breadth calculated from two phylogenetic trees (mammal supertree; maximum likelihood cytB tree) | First study to show that the proportion of zoonotic viruses per species increases with host phylogenetic proximity to humans. |
| **Albery et al. (2019)** | 250 pathogens across 4196 mammal species | Literature and database review: database originally collected by Olival et al. (2017) | Viruses | Quantitative: phylogenetic host breadth calculated from mammalian supertree | Host phylogenetic similarity and geographic overlap are strong predictors of viral sharing among mammal species. |
| **Guth et al. (2019)** | 67 zoonotic pathogens across 278 mammalian hosts (420 host-pathogen associations) | Literature and database review | Viruses | Quantitative: distance from humans on a cytochrome b phylogenetic tree | Zoonotic viruses found in animal hosts with increased host phylogenetic distance from humans correlate positively with morbidity and mortality in humans, but negatively with human transmissibility. |
| **This study** | Initially: 2,678 pathogens across 3,218 host species (13,671 host-pathogen associations).<br>Subsequently restricted to 2,595 pathogens across 2,656 vertebrate hosts (12,212 host-pathogen associations) | Literature and database review | Viruses, bacteria | Quantitative: phylogenetic host breadth calculated from host mitochondrial gene phylogeny | First study to show that the proportion of shared pathogens between vertebrate hosts (not just with respect to phylogenetic proximity to humans) decreases with increasing phylogenetic distance.<br>First study to show that multi-host bacteria infect more diverse hosts than multi-host viruses.<br>Zoonotic pathogens infect more non-human hosts than non-zoonotic pathogens. |

In calculating host range, rather than using a taxonomic proxy for host genetic similarity (e.g. broad animal taxonomic orders (Kreuder Johnson et al., 2015; McIntyre et al., 2014; Woolhouse & Gowtage-Sequeria, 2005) or mammalian host order (Han, Kramer, & Drake, 2016) — see Table 1) we use a quantitative phylogenetic distance measure. Such an approach has been used in a recent set of papers (Albery, Eskew, Ross, & Olival, 2019; Guth, Visher, Boots, & Brook, 2019; Olival et al., 2017) which all use an alignment of the mitochondrial gene cytochrome *b* to build a maximum likelihood tree, constrained to the order-level topology of the mammalian supertree (Fritz, Bininda□Emonds, & Purvis, 2009). As noted by Albery et al. (Albery et al., 2019), this mammalian supertree has limited resolution at the species tips. Here, we therefore extend this approach in two ways: (a) we use a concatenated alignment of nine mitocondrial genes rather than one, and (b) we also consider non-mammalian vertebrates (see Methods). We believe this represents the most extensive vertebrate host tree built to-date, and should give the most precise quantitative measure of the true genetic similarity between hosts.

To summarise, here we combine a systematic literature review for bacterial and viral host ranges with a mitochondrial multigene host phylogeny (Figure 1). We compiled a database of 2,595 bacteria and viruses which infect 2,656 vertebrate host species. Our quantiative analysis represents by far the most comprehensive picture of known host-pathogen associations.

**Figure 1.**
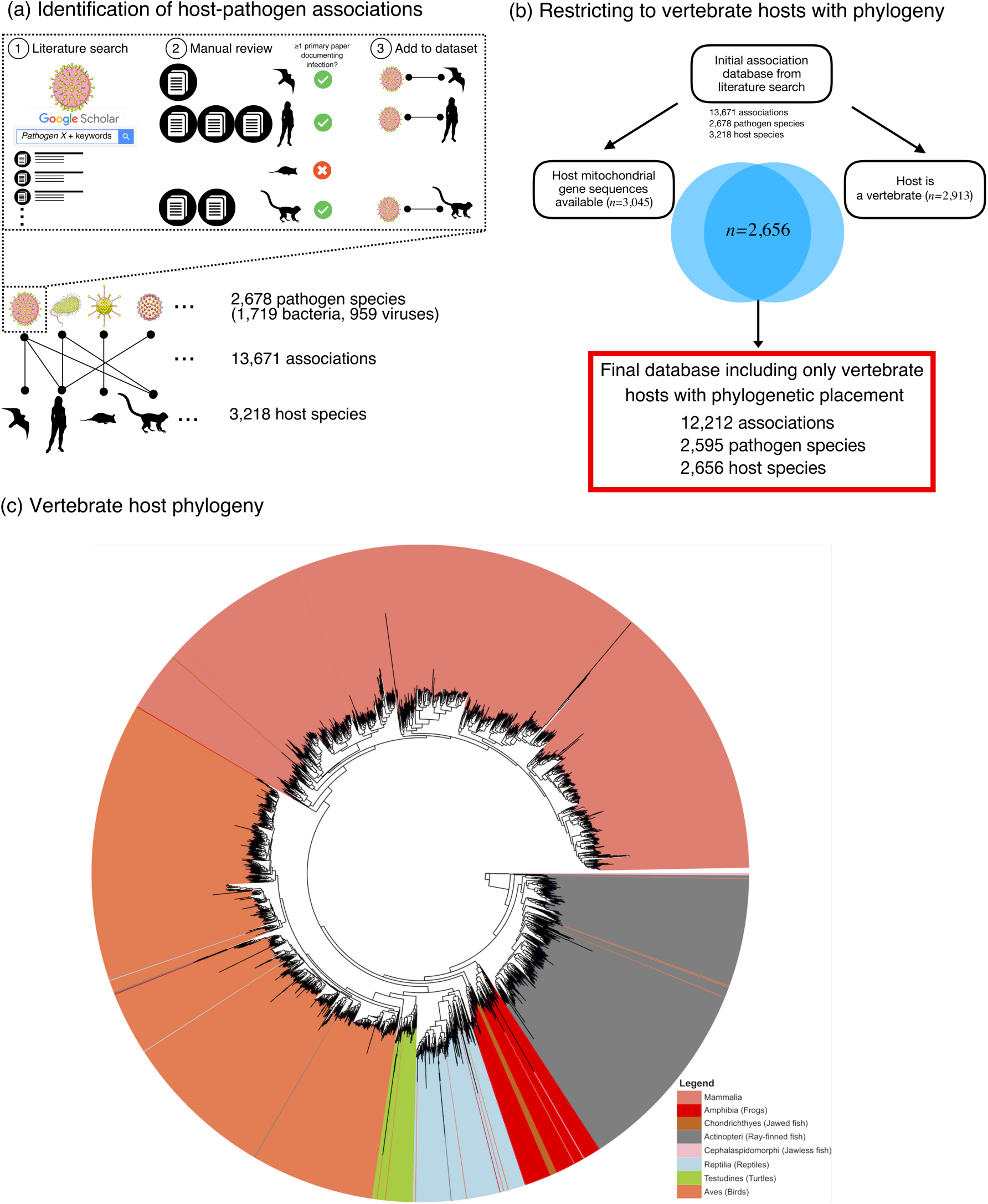
Overview of methodology for compiling the dataset. (a) Methodology of literature review (see Methods). (b) Subsetting the database to only associations involving vertebrate host species for which mitochondrial gene sequences could be identified. (c) Vertebrate host phylogeny using nine mitochondrial genes. (See ‘Image credits’ for credits and image licenses.)

## Materials and Methods

### Pathogen species

We focused on bacteria and viruses, as taken together they are the pathogen groups responsible for the majority of the burden of communicable disease in humans: the combined contribution of HIV/AIDS (viral), tuberculosis (bacterial), diarrheal diseases (predominantly bacterial and viral), lower respiratory diseases (predominantly bacterial and viral) and neonatal diseases (predominantly bacterial) made up 76.9% of all global disability-adjusted-life-years (DALYs) lost due to communicable disease in 2017 (Global Burden of Disease Collaborative Network, 2017). Bacteria and viruses also represent the two most diverse groups in terms of total number of unique pathogen species recognized (Wardeh, Risley, McIntyre, Setzkorn, & Baylis, 2015; Woolhouse & Gaunt, 2007) across both human hosts and vertebrate animals.

Bacteria and viruses infectious to humans and animals were systematically compiled by going through the complete taxonomic lists of known species from their respective authoritative organizations. Bacterial species were drawn from the LPSN 2016 release (Euzéby & Parte, 2016); and viral species were drawn from the International Committee on Taxonomy of Viruses (ICTV) 2015 release (ICTV, 2015). As such, our database is exhaustive and inclusive of all known and taxonomically recognized bacteria and virus pathogens as of December 2016. Bacteria and virus species found in abiotic environments or only in non-vertebrate hosts were not included in our database.

### Pathogen metadata

We collected further metadata for each pathogen species. Where available, we used the NCBI Genome Report for a species (last downloaded: 12^th^ March 2019) to include the mean genome size, number of genes, and GC content. We also annotated each pathogen for the presence of known invertebrate vectors (i.e. whether they can be ‘vector-borne’). For bacteria, we additionally included information on Gram stain, bacterial motility, spore formation, oxygen requirement, and cellular proliferation. These traits were collated primarily from the GIDEON Guide to Medically Important Bacteria (Berger & Berger, 2016), but where information was missing we also searched the primary literature. For viruses, we also included Baltimore classifications from the ICTV Master Species List (ICTV, 2015). For the analysis including host traits for direct comparison with Olival et al. (2017), we used their dataset of the number of disease-related publications for species (Olival et al. searched ISI Web of Knowledge and PubMed using the scientific binomial AND topic keyword: disease* OR virus* OR pathogen* OR parasit*).

### Pathogen-host interactions

Our literature search was designed to be as exhaustive and systematic as possible (Figure 1a). We used Google Scholar to conduct a literature search to verify if each bacterial or viral species was associated with a human or vertebrate animal host. Search terms consisted of the pathogen species name and the keywords: ‘infection’, ‘disease’, ‘human’, ‘animal’, ‘zoo’, ‘vet’, ‘epidemic’ or ‘epizootic’. At least one primary paper documenting the robust interaction (i.e. infection) of the bacteria or virus species with a host species needed to be found in our search for the association to be included in our database. In addition, several reputable secondary sources were used to further validate the identified pathogen-host interactions: the GIDEON Guide to Medically Important Bacteria (Berger & Berger, 2016); the Global Mammalian Parasite Database (Nunn & Altizer, 2005); and the Enhanced Infectious Diseases Database (EID2) (Wardeh et al., 2015) (eid2.liverpool.ac.uk/).

We aimed to manually read all publications found with Google Scholar searches using our keyword search terms. However, as some pathogen species are extremely well-studied and manual review of all returned publications was not possible, we decided to read only the first ten pages of search results ordered by ‘relevance’ (equivalent to a limit of 200 publications). Obviously, species with >200 results tend to be either well-studied pathogens (e.g. ‘*Mycobacterium tuberculosis*’ + ‘infection’: 62,900 results in 2016) or species with prolific host ranges (e.g. ‘*Chlamydia psittaci*’ + ‘infection’: 19,700 results in 2016). For these species we cannot claim to have captured all known hosts with our manual review; i.e. we may not have documented every single host species the pathogen has been recorded as infecting. However, we are confident that we managed to reasonably approximate the full taxonomic breadth of host range, since the first ten pages of results for these well-studied pathogens usually contained specialized review papers listing the vertebrate host species in which infections had been documented.

The majority of bacterial and viral pathogens in our database are known to cause disease symptoms in at least one of their host species. However, in order to be as comprehensive as possible, we considered as a pathogen any species for which there was *any* evidence that it can cause symptomatic adverse infections under natural transmission conditions, even if rare, including: cases where the relationship with host species is commonly asymptomatic, cases where the relationship is only symptomatic in neonatal or immunocompromised individuals, or where only a single case of infection has been recorded to date. Cases of deliberate experimental infection of host species were excluded from our database as we judged that these did not constitute natural evidence of a host-pathogen association.

A minority of bacterial and viral species in our database have not, to date, been shown to cause any infectious symptoms in the host species they naturally infect. However, characterizing symptoms in wild animal populations is difficult. Furthermore, these pathogens are often very closely related to pathogens which are definitely known to cause disease in the same hosts. For example, *Corynebacterium sphenisci* was isolated in a single study from apparently healthy wild penguins (Goyache et al., 2003) but is related to species which are pathogens across vertebrate hosts e.g. *Corynebacterium pseudotuberculosis*, the causative agent of lymphadenitis. Therefore, we included all species apart from bacteria and viruses which we considered to be clearly non-pathogens i.e. well-studied commensal or mutualistic examples such as *Lactobacilli* in the human microbiome (Walter, 2008). Important invertebrate vector species were also documented in our database, but our main analysis was restricted to vertebrate hosts.

### Host species

The taxonomic status of each host species identified in the primary literature was brought up to date by identifying the current taxonomically valid species name using the ITIS Catalog of Life (Rosokov et al., 2016) and the NCBI Taxonomy Database (www.ncbi.nlm.nih.gov/taxonomy). In some cases, hosts were not identified to the species level, but were retained in our database if they were identified to the family/order level and there were no other host species from the same family/order infected by the same pathogen species. In other cases, hosts were identified to the sub-species level (e.g. *Sus scrofa domesticus*) if these sub-groups were economically and/or sociologically relevant.

The full compiled database contained 13,671 associations (Figure 1a), including invertebrate hosts (*n*=305) as well as vertebrates (*n*=2,913). However, we restricted our host-relatedness analysis to vertebrates for which we could construct a mitochondrial gene phylogeny (Figure 1b).

### Definition of zoonosis

We classified a pathogen as zoonotic if it infected both humans and additional vertebrate animals, including those shared but not known to be naturally transmissible among different host species, unlike the WHO’s definition of ‘any disease or infection that is naturally transmissible from vertebrate animals to humans and vice-versa’ (WHO, 2019). This definition includes species that mostly infect their various hosts endogenously or via the environment (i.e. opportunistic pathogens) such as species in the bacterial genus *Actinomyces*. We did this based on the observation that many new infectious diseases occur through cross-species transmissions and subsequent evolutionary adaptation. Pathogens could also evolve to become transmissible between host species. In addition, we are interested in how the overall host range and host relatedness of a pathogen effects its likelihood of emergence and its association with other pathogen characteristics. We did not classify a bacterial or viral species as zoonotic if it had only been recognized outside of human infection in invertebrate hosts.

### Host phylogeny

To infer a phylogenetic tree for all 3,218 vertebrate and invertebrate species, we relied on nine mitochondrial genes: *cox2, cytb, nd3, 12s, 16s, nd2, co3, coi*, and *nadh4*. Our strategy was as follows. First, we collected mitochondrial genes for species that had mitochondrial gene submissions present in the NCBI database. For species without a mitochondrial gene submission but where a whole genome was present, we extracted the genes by blasting the genes of a taxonomically closely related species and then extracting the gene from the resulting alignment. If no mitochondrial gene or whole genome submissions were available, we used the NCBI taxonomy to approximate the species using a closely related species (using either available genes or sequences extracted from genomes). Using this strategy and some manual filtering, we were able to obtain mitochondrial gene sequences for 3,069 species (including invertebrates).

We merged these genes in their distinct orthologous groups (OGs) using OMA (Altenhoff et al., 2018). We used the nine largest OGs that had our expected nine genes as a basis for alignment to ensure that alignment was conducted on high quality related sequences. We aligned sequences for each OG separately using mafft (v7) with the options ‘--localpair -- maxiterate 1000’ (Katoh & Standley, 2013). We then used MaxAlign (v1.1) (Gouveia-Oliveira, Sackett, & Pedersen, 2007) to get the best aligning sequences from all sequences. In order to produce more consistent alignment when only partial gene submissions were available, we used the ‘--add’ parameter of mafft to append all the residual sequences that were part of a corresponding OG. Then, we concatenated all OGs and inferred the phylogenetic tree using IQ-TREE (v1.5.5) with the options ‘-bb 1000’ and the HKY+R10 model as identified by ModelFinder part of the IQTREE run (Hoang, Chernomor, von Haeseler, Minh, & Vinh, 2018; Nguyen, Schmidt, von Haeseler, & Minh, 2015).

We observed a strong similarity between cophenetic distances for the 551 mammals in our multi-mitochondrial gene tree that are also included in the *cytb* tree produced by Olival et al. (Olival et al., 2017) and used by subsequent studies (Albery et al., 2019; Guth et al., 2019), but there were some discrepancies (Supplementary Figure 1). We did not investigate these further, but suspect they may have stemmed from us not constraining our phylogeny to an existing order-level topology; that is, our phylogeny represents host genetic distances inferred solely from available mitochondrial gene sequences, agnostic to any other information. However, this tree appeared to be globally highly consistent with NCBI taxonomic ordering, with only a small minority of species disrupting monophyly of groups (*n=*93, 3.1%). The apparently incorrect placement of these species could have several possible explanations, including: mislabelling in the database, poor sequence quality, or problems with the tree inference. After pruning the tree to include only vertebrate species (*n*=2,656, Figure 1c), a reduced fraction of species disrupted monophyly of groups (*n*=40, 1.5%). The analyses presented in the main text include species which disrupted monophyly.

### Phylogenetic host breadth

Following Olival et al. (Olival et al., 2017) we define the Phylogenetic Host Breadth (PHB) of a pathogen as a function (*F*) of the cophenetic matrix of pairwise distances *d*_*ij*_ between its *N* hosts. Specifically, we take the function over the upper triangle of this (symmetric) matrix:

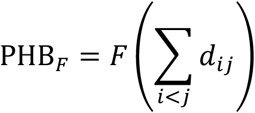

We found that the mean PHB for a pathogen (i.e. the average inter-host distance) was correlated with the maximum PHB (i.e. the largest inter-host distance) (Supplementary Figure 2), so there was little practical difference in using either measure. For simplicity, PHB refers to PHB_mean_ unless otherwise stated. In fitting generalized additive models to predict zoonotic potential, models included different PHB quantities (see below).

### Statistical analyses

Code for all analyses is available in our code and data repository (https://github.com/liampshaw/Pathogen-host-range). Here, we give a brief overview of our statistical methods.

#### Descriptive analyses of pathogen traits and host range

To summarise the dataset and make the most use of the traits we manually compiled, we performed separate analyses of different pathogen traits and their associations with host range using a specialist/generalist distinction. We used Chi-squared tests to assess whether viruses and bacteria differed in their proportion of specialist pathogens or vector-borne pathogens. We used Wilcoxon rank sum tests to compare the distributions of GC content and genome size for specialists and generalists within viruses and bacteria. We used Chi-squared tests to compare the distribution proportion of generalists within subsets of bacteria based on lifestyle factors: motility, cellular proliferation, spore formation, and oxygen requirement. Reported p-values are not corrected for multiple testing. These results are best viewed as exploratory; some of the conclusions were not retained when looking at the partial effects of predictors in a best-fit GAM, controlling for the effects of other variables.

#### Generalized additive models (GAMs)

We fitted GAMs to rank predictors using the approach of Olival et al. (2017), adapting their previously published code. Their approach uses smoothed spline predictors and automated double penalty smoothing to eliminate redundant terms from the full model (select=TRUE in ‘gam’ function). Where multiple variables measured the same effect, alternate GAMs used only one of these variables (e.g. different proxies for research effort, although in practice these metrics are highly correlated). Categorical and binary variables are fitted as random effects for each level of the variable.

#### Host traits associated with greater pathogen richness

We used a dataset of host traits for terrestrial wild mammal species compiled by Olival et al. (2017) to predict pathogen richness (bacterial and viral) per species. These host traits included a phylogenetic eigenvector regression (PVR) of body mass. As Olival et al. collected information for an analysis of only viral pathogens, we found there was better overlap for viruses in our dataset (*n*=613 host wild mammals) than for bacteria (*n*=274).

#### Zoonotic potential

For bacteria, GAMs could include terms for host range (PHB_mean_, PHB_median_, or PHB_max_), research effort (NCBI PubMed, Nucleotide, or SRA results), motility, sporulation, being vector-borne, oxygen requirements, and Gram stain. We excluded cellular lifestyle (intra/extracellular) as a predictor due to low numbers, and excluded pathogens of unknown motility (*n*=50) or sporulation (*n*=17). For viruses, GAMs could include terms for host range, research effort, genome size (number of proteins and length), being vector-borne, and genome type (Baltimore classification). We excluded pathogens with unknown genome size (*n*=253). We observed structure in some partial effect residuals in the best-fit GAMs: research effort for bacteria (Figure 6b) and host range for viruses (Figure 6d). This structure was driven by pathogen taxonomy, with families (orders) for bacteria (viruses) having different zoonotic potential; e.g. the *Staphylococcaceae* contain a high proportion of generalists. Attempts to include taxonomy as a categorical predictor produced best fit models which excluded all lifestyle factors (not shown), although host range and research effort were still the strongest predictors.

**Figure 2.**
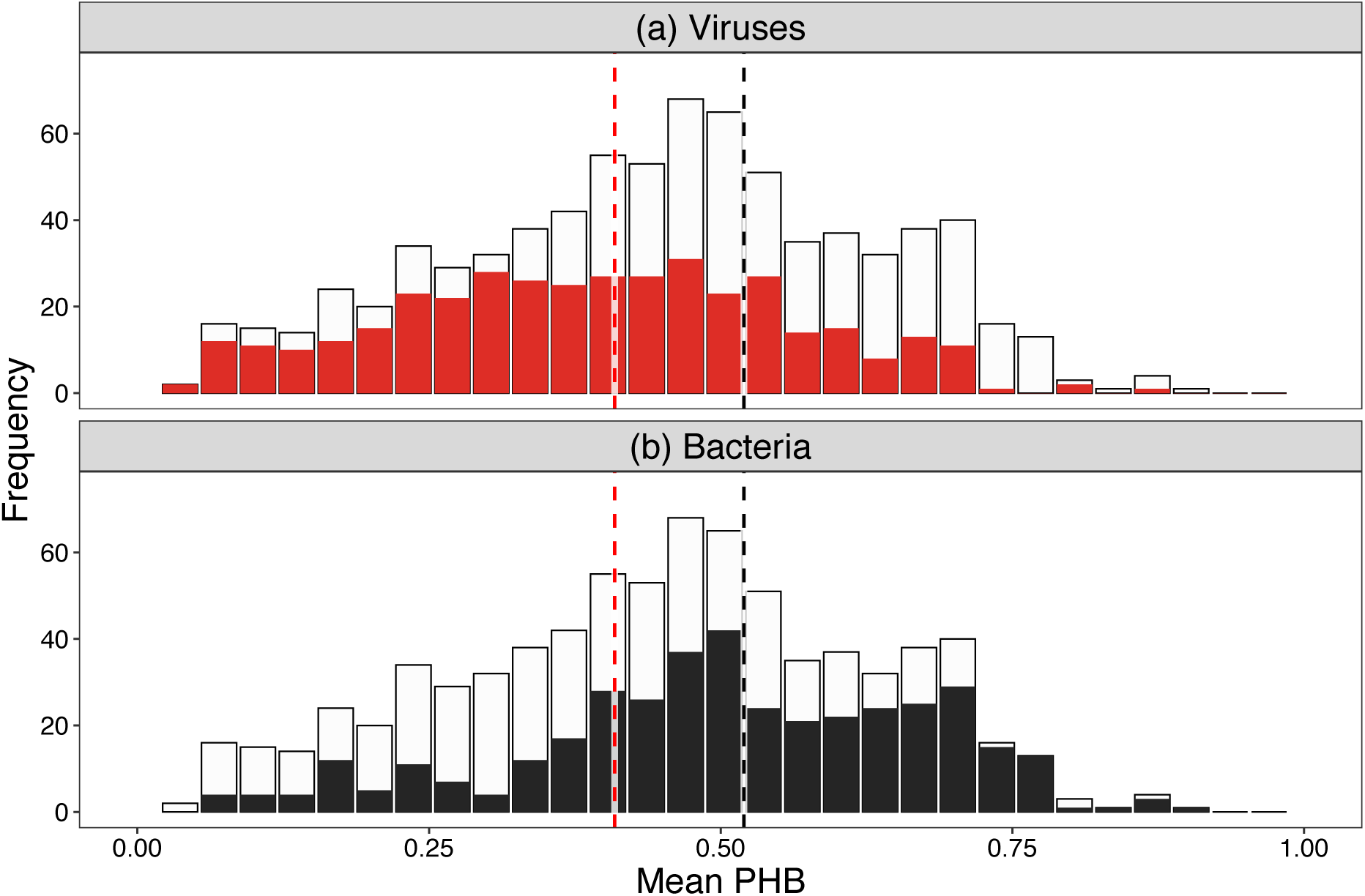
Mean phylogenetic host breadth for multi-host (top panel) bacteria and (bottom panel) viruses. Bacteria and viruses are shown in black and red respectively, with the overall pathogen histogram (both types) shown in grey on both panels to help comparison. On average, multi-host bacteria have a more diverse host range than viruses (black/red dashed lines indicate median for bacteria/virus respectively). The majority of pathogens have a mean PHB < 0.03 (*n*=1,816, 70.0%) and are excluded from the plot.

**Figure 3.**
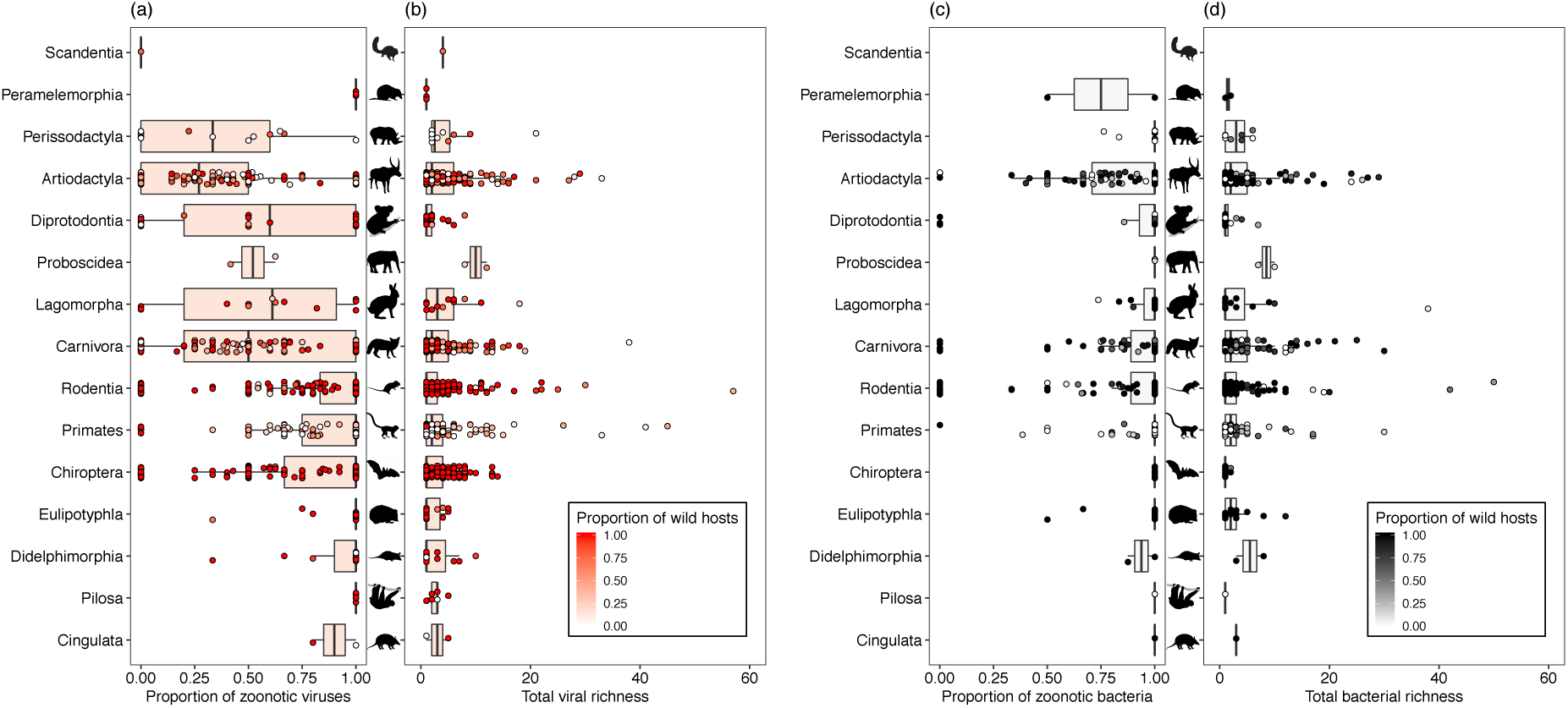
Observed pathogen richness in mammals. Box plots of (a,c) proportion of zoonotic viruses/bacteria and (b,d) total viral/bacterial richness per species, aggregated by order. Each point represents a host species, with the colour indicating the proportion of associations for a host species derived from observations of wild hosts (as opposed to in domestic or captive hosts of the same species). Lines represent median, boxes, interquartile range for each order. Data from 3,869 host–virus associations and 2,653 host-bacteria associations. The ordering of mammals is the same as in Figure 1 of Olival et al. (2017) to facilitate comparison. Richness plots have a cutoff at 60 to increase clarity of boxplots, so a small number of outlier species with greater pathogen richness are not shown (outliers for viral richness: one in Perissodactyla, four in Artiodactlya; outliers for bacterial richness: one in Perissodactyla, four in Artiodactyla, one in Carnivora). (See ‘Image credits’ for credits and image licenses.)

**Figure 4.**
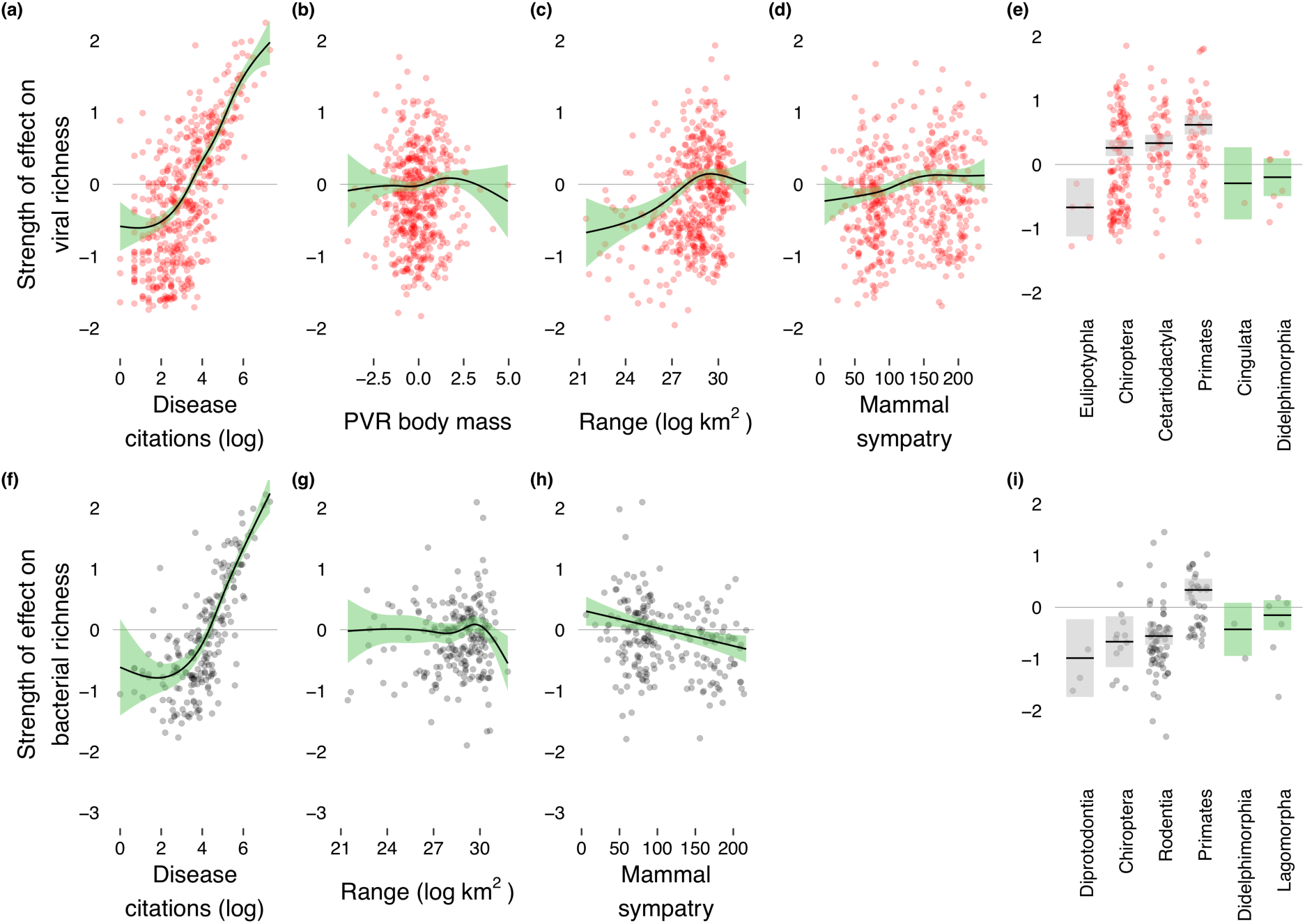
Host traits which predict total viral richness (top row) and bacterial richness (bottom row) per wild mammal species. Each plot shows the relative effect of the variable in the best-fit GAM accounting for the effects of other variables (see Table 3 for numerical values). Shaded circles represent partial residuals and shaded areas represent 95% confidence intervals around the mean partial effect. (a, f) Number of disease-related citations per host species, (b) phylogenetic eigenvector representation (PVR) of body mass i.e. corrected for phylogenetic signal, (c, g) geographic range area of host species, (d, h) number of sympatric mammal species overlapping with at least 20% area of target species range, and (e, i) mammalian order (non-significant terms retained in best model shown in grey). For bacteria, PVR body mass was not included in the best-fit GAM.

**Figure 5.**
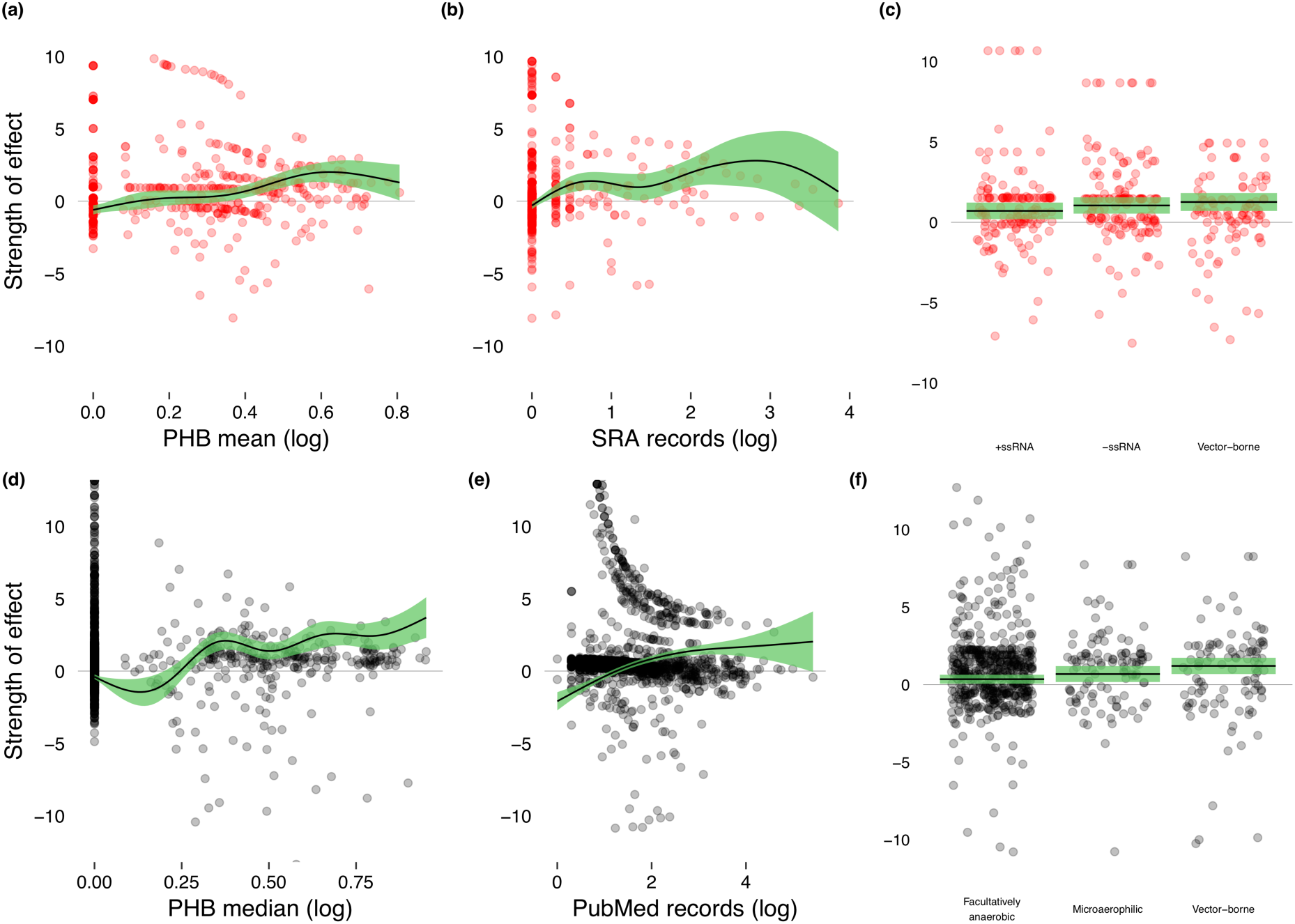
The partial effects of predictors in the best-fit GAM for predicting whether a bacterium (top row) or a virus (bottom row) is zoonotic. Each plot shows the relative effect of the variable in the best-fit GAM accounting for the effects of other variables (see Table 3 for numerical values). Shaded circles represent partial residuals and shaded areas represent 95% confidence intervals around the mean partial effect. Viruses: (a) Mean phylogenetic breadth of viral pathogen, (b) SRA records for viral pathogen, (c) Significant categorical predictors. +ssRNA and -ssRNA are mutually exclusive as they come from the ‘genome type’ variable. Bacteria: (d) Median phylogenetic breadth of bacterial pathogen, (e) PubMed records for bacterial pathogen, (f) Significant categorical predictors. Facultatively anaerobic and microaerophilic are mutually exclusive as they come from the ‘oxygen’ lifestyle variable. Gram stain and motility were included in the best-fit model but were not significant. Predictors are different for each best-fit GAM because the model term for e.g. phylogenetic breadth could be chosen from the mean, median, or maximum PHB for a pathogen.

**Figure 6.**
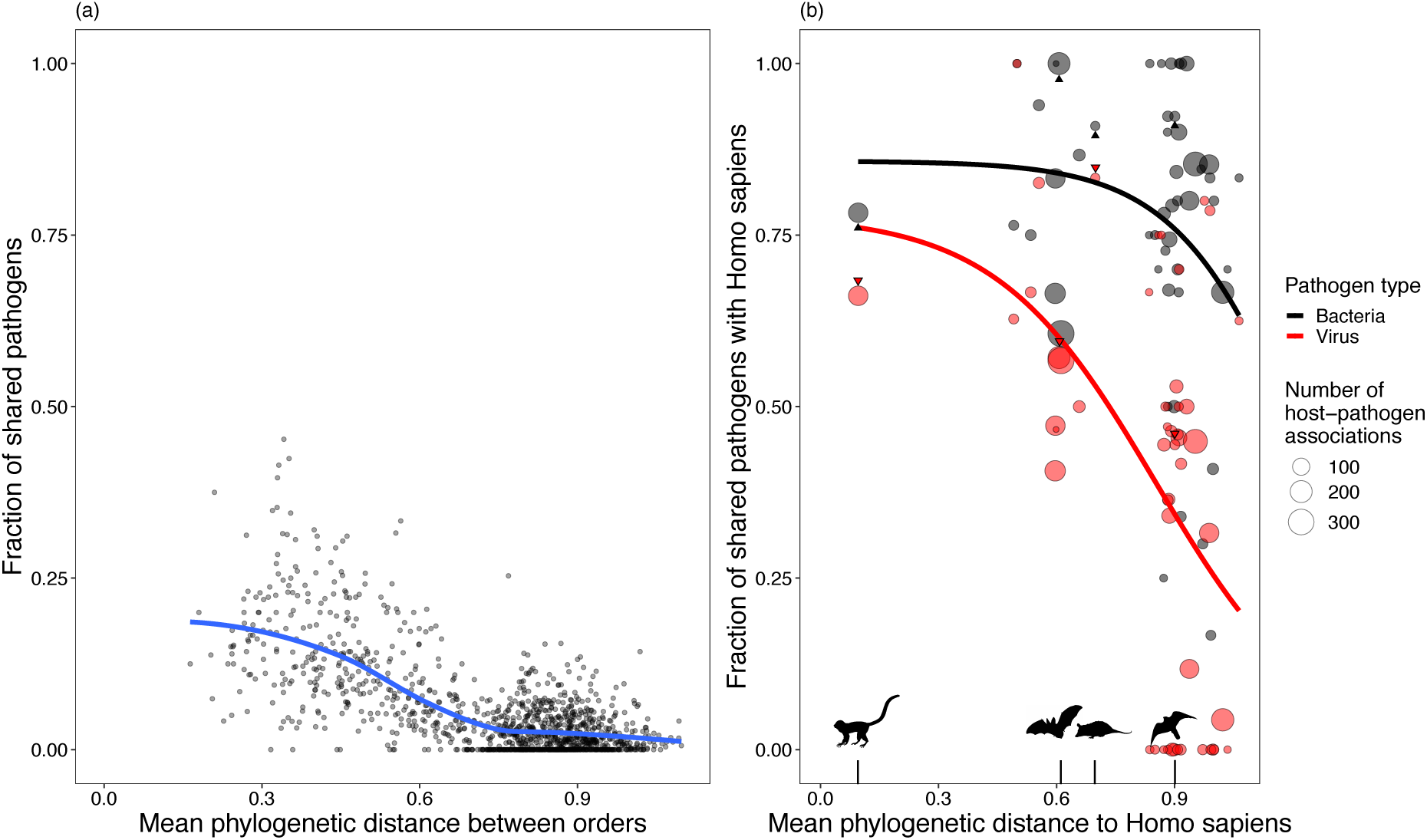
The fraction of shared pathogens between hosts decreases with inter-host phylogenetic distance. For the definitions of fraction of shared pathogens between two taxa, see Methods. (a) All pairwise comparisons between different host orders with at least 10 host-pathogen associations in the database. The blue line shows a smoothed average fit, produced with ‘loess’. Only comparisons between host orders with at least 10 host-pathogen associations in the database are shown. (b) Fraction of shared pathogens between these host orders and *Homo sapiens* (as a fraction of total unique pathogens infecting at least one species in each order), showing data for bacteria (black) and viruses (red) together with a sigmoidal fit (thick line) for each pathogen type. Size of points indicates the number of unique host-pathogen associations for that order. Four illustrative orders are indicated with silhouettes (Primates, Chiroptera, Didelphimorphia, Falconiformes) with black (red) arrows indicating the relevant point for bacteria (viruses). (See ‘Image credits’ for credits and image licenses.)

#### Shared pathogen analysis

If we denote the set of pathogens seen at least once in a host taxon *a* as *p*_*a*_ (where the taxon could be a species, genus, family etc.), we define the fraction of shared pathogens between two taxa *a* and *b* as:

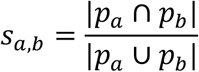

Note that this definition is symmetric in *a, b.* It can therefore be compared with the (mean) phylogenetic distance between taxa using a Mantel test to determine the correlation.

Another property we consider is the fraction of the pathogens seen in a host taxon which are also seen in a reference host species (e.g. humans). Taking the comparison of primates and humans as an example: in the database, the primates (excluding humans) are represented by 147 host species with 762 host-pathogen associations. The total number of pathogen species with at least one association with a primate species is *p*_P_=222. Of these, 158 are also seen in humans (the total number of pathogen species with at least one association with humans is *p*_H_=1,675). We can then define the fraction of pathogens of primates also seen in humans as

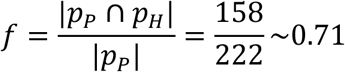

We used an illustrative sigmoidal fit of the form 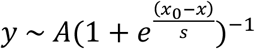 separately for bacteria and viruses to model the relationship between *f* and the mean phylogenetic distance of the order to humans.

## Results

### A comprehensive database of pathogen associations for vertebrates

Our database includes 12,212 associations between 2,595 vertebrate pathogens (1,685 bacterial species across 127 families; 910 viral species across 35 families) infecting 2,656 host species across 90 host orders. Pathogens infecting *Homo sapiens* made up 1,675 of all associations (13.7%), the largest single host species group. The viral and bacterial pathogens with the most recorded host associations were Newcastle disease virus (NDV) (*n*=207, 1.7%) and *Chlamydia psittaci* (*n*=133, 1.1%), respectively.

### Specialist pathogens are the most common category

Approximately half of all pathogens infected only a single host species (*n=*1,473, 56.8%; Table 2). For pathogens not infecting humans, specialists were less common than generalists. Bacteria had a significantly higher proportion of specialists compared to viruses (64.5% vs. 42.5%; *χ*^2^ test: *χ*^2^[df=1, *n*=1,635]=5.77, *p*=0.016). Almost half of all bacteria were human specialists (855 of 1,685, 50.7%). Despite the dominance of specialists, many generalist pathogens had broad host ranges spanning more than one host order: around one in three pathogens infected multiple host orders (*n=*508, 30.3% of bacteria, *n*=307, 33.7% of viruses; Table 2).

**Table 2.**
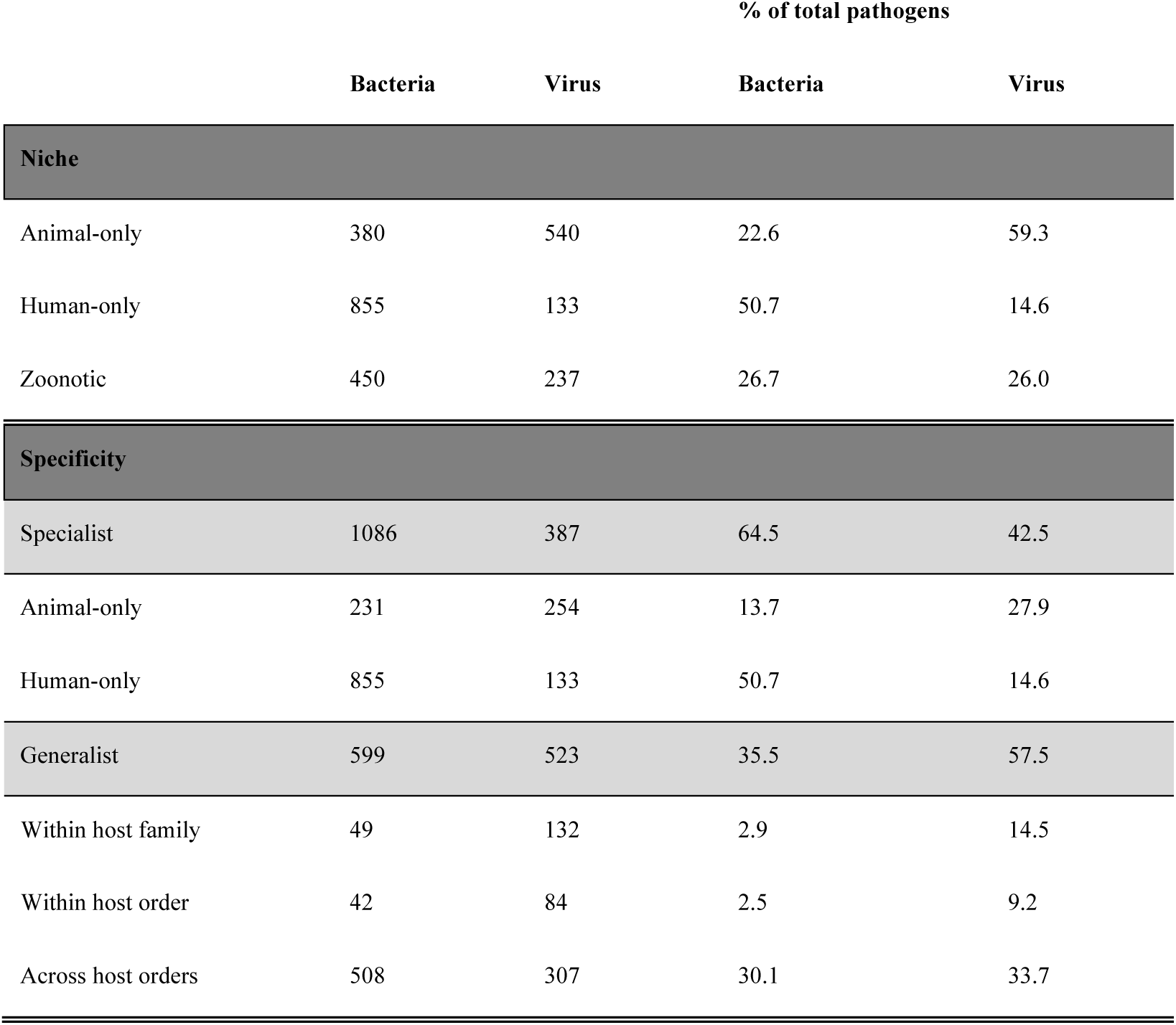
Summary of pathogen niche and specificity types. A ‘specialist’ pathogen infects only a single host, a ‘generalist’ more than one. Generalists are categorised according to whether their hosts are within the same family (e.g. Bovidae), order (e.g. Artiodactyla), or across orders. Percentages are of the total pathogen species of each type (bacteria or virus).

Considering well-represented pathogen taxonomic families (>20 pathogens in association database), the bacterial family with the highest proportion of generalists was *Staphylococcaceae* (24 of 29, 83%; Supplementary Figure 3). For viruses, it was *Bunyaviridae* (68 of 79, 86%; Supplementary Figure 4).

### Multi-host viruses have a more restricted host range than multi-host bacteria

Although the majority of pathogens infect just one host, and the total proportion of bacteria and viruses infecting multiple host orders was similar (30.1% vs. 33.7%), the distribution of generalists was significantly different between bacteria and viruses. Multi-host viruses were more likely than bacteria to only infect a single host family (Table 2). A minority of pathogens were vector-borne (*n*=272, 10.5%), and viruses were significantly more likely to be vector-borne than bacteria (18.4% vs. 6.2%; *χ*^2^ test: *χ*^2^[df=1, *n*=2,589]=91.5, *p*<0.001). A higher proportion of vector-borne viruses were generalists than those which were not vector-borne (70.1% vs. 36.6%; *χ*^2^ test: *χ*^2^[df=1, *n*=907]=60.9, *p*<0.001). The same was true for bacteria (49.5% vs. 21.4%; *χ*^2^ test: *χ*^2^[df=1, *n*=1,682]=42.3, *p*<0.001) (Supplementary Table 1).

This restricted host range of multi-host viruses was also apparent in the distribution of mean phylogenetic host breadth (PHB) for multi-host pathogens (Figure 2). Bacteria generally had a more positively skewed distribution of mean PHB compared to viruses (Figure 2; median 0.520 vs. 0.409, *p*<0.001 Wilcoxon rank sum test). Notably, these distributions were both above the median maximum phylogenetic distance between hosts from the same order, which was 0.323. The observation that bacteria had a more positively skewed distribution of mean PHB was reproduced when subsampling to exclude human hosts for both domestic and non-domestic hosts (see repository).

### Pathogen richness varies by host order

Observed pathogen richness varied at the level of host order (Figure 3). Considering only host species with an association with at least one bacterial species and one viral species, bacterial and viral richness were strongly correlated (Spearman’s *ρ*=0.57, *p*<0.001). The proportions of these bacteria and viruses shared with humans were more weakly correlated (Spearman’s *ρ*=0.21, *p*<0.001).

We used a dataset of host traits for wild mammals previously compiled by Olival et al. (2017) to find predictors of total bacterial and viral richness within a species using GAMs (Figure 4). More than 60% of total deviance was explained by the best-fit GAMs (Table 3a and 3b) for bacteria and viruses respectively). The number of disease-related citations for a host species was the strongest predictor of the number of both bacterial and viral pathogens, accounting for ∼80% of relative deviance.

**Table 3.**
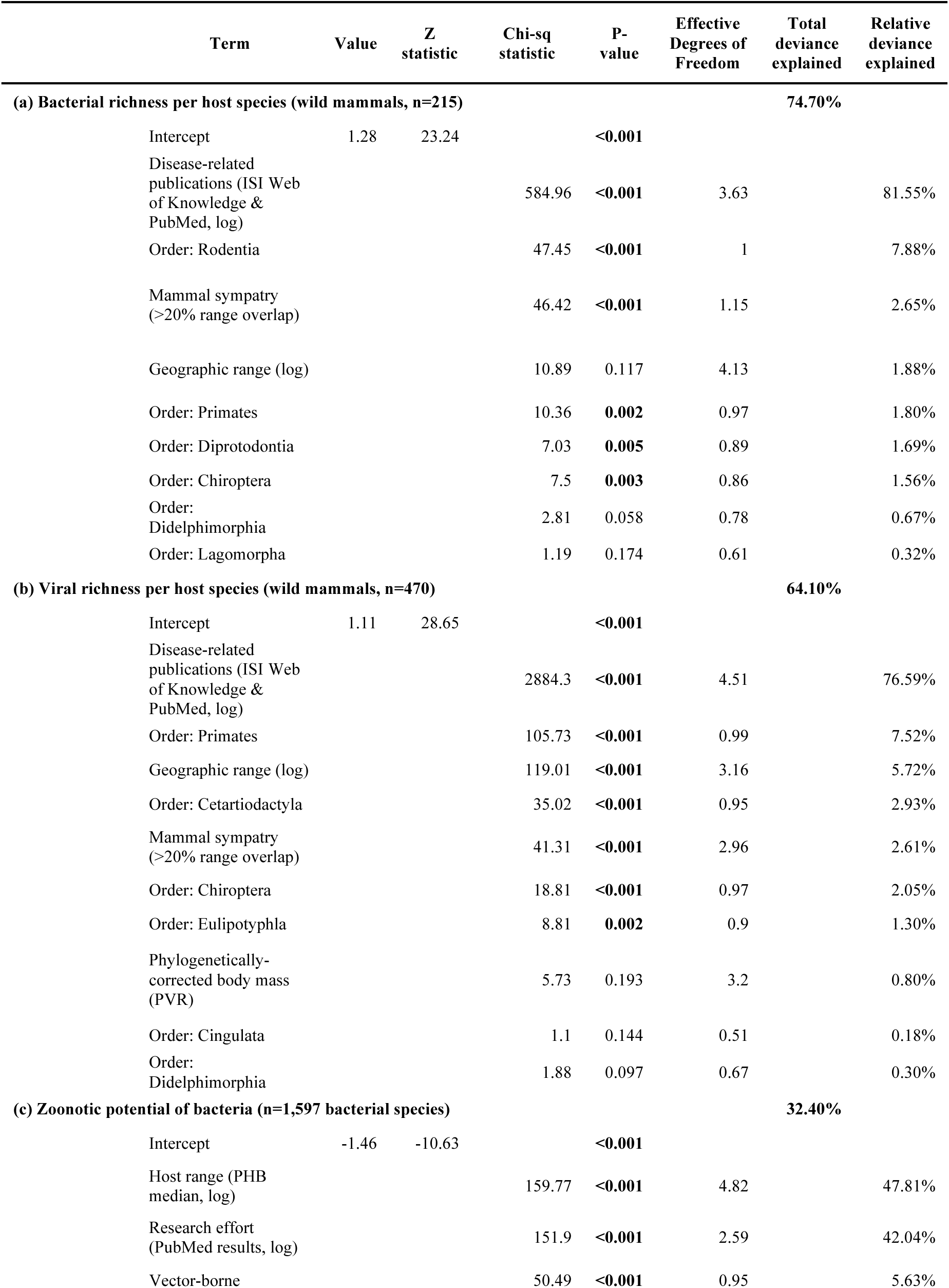
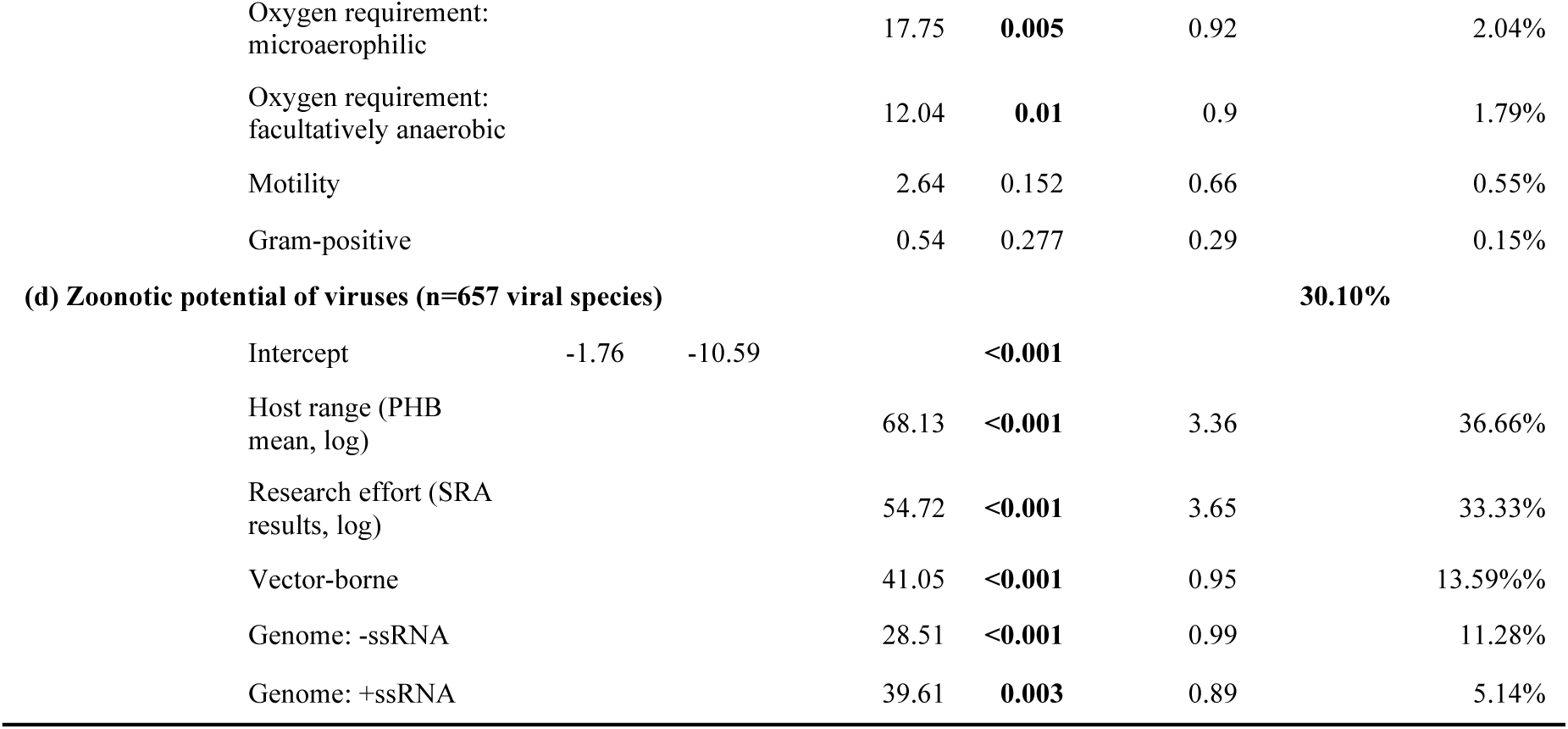
Summary of best-fit GAMs for predicting bacterial or viral richness per wild mammal species, and the probability of a pathogen being zoonotic.

### Pathogen genome and host range

We observed different distributions of pathogen genome GC content and genome size depending on whether a pathogen was a specialist or a generalist (Supplementary Figure 5). We had information on the number of proteins for *n=*657 viruses (72.2%, 5,815 associations). While there was no significant correlation between the number of proteins in a virus genome and mean PHB (Spearman’s *ρ*=0.06, *p*=0.13), there was a significant positive correlation between genome size and PHB (Spearman’s *ρ*=0.23, *p*<0.001).

We had information on genome GC content and genome size for a similar proportion of bacteria (*n*=1,135, 67.4%, 4,619 associations; *n=*550 species lacked data). However, there was no significant correlation between bacterial genome size and PHB (Spearman’s *ρ*=-0.05, *p*=0.10), although specialists had a slightly larger genome size than generalists (means: 3.66 vs. 3.30 Mb; Wilcoxon rank sum test: *W*=140,610, *p*=0.007).

### Pathogen factors affecting host range of viruses

#### Genome composition

Viruses with RNA genomes had a greater PHB than DNA viruses (median: 0.238 vs. 0). Subsetting further, +ve-sense single-stranded RNA viruses (Baltimore group V) had the greatest PHB (Supplementary Figure 6).

DNA viruses typically have much larger genomes than RNA viruses. We therefore fitted a linear model for mean PHB using both DNA/RNA genome and genome size, with an interaction term. Having an RNA genome and a larger genome were both significantly associated with greater mean PHB in this linear model (*t*=6.11 and *t*=4.58 respectively, *p*<0.001 for both, Supplementary Table 2), with a non-significant interaction between them (*p*=0.36). In line with this, we found that the proportion of zoonotic viruses was higher for RNA viruses (198 of 572, 34.6%) than DNA viruses (33 of 286, 11.5%), in agreement with Olival et al. (2017) who found a similar proportion (41.6% vs. 14.1%).

### Pathogen factors affecting host range of bacteria

We looked at the effect of bacterial lifestyle factors on the proportion of specialist and generalist pathogens (Figure 7).

#### Motility

The majority of bacterial species in our database were non-motile (non-motile: *n=*1,121, motile: *n*=514, not applicable: *n=*50 e.g. *Mycoplasmatales*). Motile bacteria were more likely to infect multiple hosts compared to non-motile bacteria (27.2% vs. 21.7%; *χ*^2^ test: *χ*^2^[df=1, *n*=1,635]=5.77, *p*=0.016).

#### Cellular proliferation

Bacteria with an extracellular lifestyle (*n=*161) were not more likely to infect multiple hosts compared to obligate (*n*=53) or facultative (*n*=93) intracellular pathogens (*χ*^2^ test: *χ*^2^[df=2, *n*=307]=0.29, *p*=0.86). Combining motility and cellular proliferation in a linear model suggested that neither was associated with greater mean PHB (Supplementary Table 3).

#### Spore formation

Only a small number of bacterial pathogens were spore-forming (*n*=91), and they did not have a significantly different number of generalists compared to non-spore-forming bacteria.

#### Oxygen requirement

The oxygen requirements of bacteria were significantly associated with the proportion of generalists in each group (*χ*^2^ test: *χ*^2^[df=2, *n*=1,573]=55.5, *p*<0.001). Aerobic bacteria (*n*=648) were nearly twice as likely to infect multiple hosts compared to anaerobic bacteria (*n*=343) (20.8% vs. 10.8%). However, facultatively anaerobic bacteria (*n*=582) had an even higher proportion of species infecting multiple hosts (31.5%).

### Predicting zoonotic potential from pathogen traits

We fitted GAMs to predict whether or not a pathogen was zoonotic using pathogen traits and inspected the partial effects for each predictor in the best-fit model (Figure 5). Best-fit GAMs could explain ∼30% of total deviance (Table 3). We found that research effort and host range (excluding human hosts) were the two strongest predictors of zoonotic potential, together accounting for >70% of relative deviance. For bacteria, being facultatively anaerobic or microaerophilic were significantly associated with zoonotic potential (Figure 5c, Table 3c) ; for viruses, those with an RNA genome had greater zoonotic potential (Figure 5f, Table 3d). Vector-borne pathogens had greater zoonotic potential for both bacteria and viruses.

### Pathogen sharing between hosts decreases with phylogenetic distance

The proportion of total pathogens shared between host orders decreased with phylogenetic distance (Figure 6a). Comparing vertebrate host orders specifically to *Homo sapiens* showed that the closer an order was to humans, the greater the fraction of pathogens that were shared for both bacteria and viruses, with an approximately sigmoidal relationship (Figure 6b). The decrease in the fraction of shared pathogens was steeper for viruses than bacteria.

## Discussion

In this work, we have compiled the largest human-curated database of bacterial and viral pathogens of vertebrates across 90 host orders. To date, this represents the most detailed and taxonomically diverse characterization of pathogen host range. Using this database, we have been able to conduct a detailed quantitative analysis of the overall distribution of host range (host plasticity) across two major pathogen classes (together bacteria and viruses comprise the majority of infectious diseases). We also use this database to examine the proportion of pathogens shared among host orders.

We found that pathogen sharing was strongly correlated with the phylogenetic relatedness of vertebrate hosts. This finding corroborates and generalises the observation by Olival et al. (2017) for viral pathogens of mammalian hosts, as well as other studies using smaller taxon-specific datasets (13, 15, 21). This suggests that phylogeny is a useful general predictor for determining the ‘spillover risk’ (i.e. the risk of cross-species pathogen transmission) of different pathogens into novel host species for both bacteria and viruses. Given the difficulty in predicting the susceptibility of cross-species spillovers (Parrish et al., 2008) and the restriction of most previous work to viruses, this finding is an important step in our understanding of the broad factors underlying and limiting pathogen host ranges.

The underlying mechanisms by which phylogeny affects spillover risk still need to be more closely examined. Pathogens are likely to be adapted to particular host physiologies (e.g. host cell receptors and binding sites), which are expected to be more similar between genetically closer host species. One mechanism by which a pathogen may be able to establish a broader host range is by exploiting more evolutionarily conserved domains of immune responses rather than immune pathways with high host species specificity. Such an association has been shown among viruses for which the cell receptor is known (Woolhouse, 2002). Interestingly, we found that the decrease in the fraction of shared pathogens with increasing phylogenetic distance was steeper for viruses than bacteria, which suggests that bacterial pathogens, on average, have higher host plasticity than viruses (i.e. a greater ability to infect a more taxonomically diverse host range). Future studies could examine whether host cell receptors for bacterial pathogens are more phylogenetically conserved compared with host cell receptors for viral pathogens.

When examining the overall distribution of host ranges, we found a substantial fraction of both bacterial and viral pathogens that have broad host ranges, encompassing more than one vertebrate host order. The evolutionary selection of pathogens that have broad host ranges has been a key hypothesis underpinning the emergence of new zoonotic diseases (Cleaveland et al., 2001; Woolhouse & Gowtage-Sequeria, 2005), and mean PHB has previously been shown to be the strongest predictor of the zoonotic potential of viral pathogens (Olival et al., 2017). High pathogen host plasticity has also been found to be associated with both an increased likelihood of secondary human-to human transmissibility and broader geographic spread (Kreuder Johnson et al., 2015), both of which are traits linked to higher pandemic potential. Given these observations, it may be useful to more closely monitor those pathogens with the highest mean PHBs that have not yet been identified as zoonoses.

Several traits were found to be significantly associated with bacterial and viral host ranges. For viruses, RNA viruses and larger genome size were independently associated with a broader host range. This is in line with RNA viruses appearing particularly prone to infecting new hosts and causing emerging diseases, something which has been attributed to their high mutation rate (Holmes, 2010). The positive association between viral genome size and host range might be due to pathogens specialising on a narrower range of hosts requiring a smaller number of genes to fulfil their replication cycle.

For bacteria, motile and aerobic pathogens had a wider host range, with the largest number of hosts found for facultative anaerobes, perhaps suggesting a greater ability to survive both inside and outside hosts. Conversely, we did not find a strong association between genome size and host range in bacteria; in fact, specialists had slightly larger genomes on average compared to generalists. Since genome reduction through loss of genes is a well-recognised signature of higher virulence in bacteria (Weinert & Welch, 2017), this suggest that pathogenicity may be largely uncorrelated to host range in bacteria. It would be interesting to further explore these relationships for obligate and facultative pathogens in the future.

We found a surprising lack of association between the expected ‘intimacy’ of host-pathogen relationships (as judged with pathogen lifestyle factors) and host range. We identified more single-host bacteria than viruses, which was the opposite of what we predicted going into this study. One possibility is that bacterial pathogens may be more dependent on the host microbiome i.e. their ability to infect other host species may be more contingent on the existing microbial community, compared to viruses. However, we recognize that literature bias could contribute to this conclusion, particularly for RNA viruses which are more difficult to identify and diagnose than other infective agents. We also found that intra-cellular and extra-cellular bacteria had roughly the same number of hosts despite our expectation for intra-cellular bacteria to have a more narrow host range due to their higher expected intimacy with their host. However, it should be noted that information about cellular proliferation was only available for 18% (307 of 1,685) of all bacteria in the database, and this is a trait which can be difficult to unambiguously characterize (Silva, 2012).

Previous studies of viral pathogens have shown that those that are vector-borne tend to have greater host ranges — whether through higher host plasticity (Kreuder Johnson et al., 2015) or higher mean PHB (Olival et al., 2017). We replicate this observation for both viruses and bacteria, suggesting a strong and consistent effect of being vector-borne for a pathogen. We also found that greater host range was associated with greater zoonotic potential for viral and bacterial pathogens, complementing previous work restricted to viruses (Olival et al., 2017).

We controlled for research effort (number of publications, or number of SRA records) and found that it was a strong predictor of both greater pathogen richness within a host species, and the zoonotic potential of a pathogen. However, disentangling these factors is difficult. There could be increased research efforts to study known zoonoses to identify them in animals in order to establish possible ‘reservoirs’, giving a biased picture. However, this could also partly be a consequence of the global distribution of humans and their propensity to transmit pathogens to both wild and domestic species. There are multiple documented cases of zoonotic pathogens having transmitted from humans to other animals, rather than the other way around. Prominent examples include the ancestor of the agent of tuberculosis, which humans likely transmitted to cows (Brosch et al., 2002; Mostowy et al. 2002), or the multiple host jumps of *Staphylococcus aureus* from humans to cattle, poultry and rabbits (Viana et al., 2015; Weinert et al., 2012). Such host jumps from humans to animals may contribute to the pattern of zoonotic species having broader host ranges in particular for pathogens at high prevalence in humans.

Our results have several further limitations. Our database was compiled from a comprehensive synthesis of the available evidence in the literature about host-pathogen associations. Our results are therefore necessarily biased by differences in research intensity among both different pathogen and host species; or, viewed another way, they are a fair reflection of the current state of knowledge in the literature. For example, specialist pathogens of humans were the largest single group of bacterial species most likely because these have been comparatively well-studied. Research effort targeted towards a pathogen primarily reflects its medical and/or economic impact, and as such does not invalidate a classification into specialist and generalist pathogens. We do not expect research effort to *a priori* bias comparisons of host range between viruses and bacteria, or subsets therein defined by particular traits such as GC content. Although we attempted to control for research effort in our statistical analyses which did not depend on a binary ‘specialist’ vs. ‘generalist’ distinction, the limitation of reflecting the current state of knowledge still applies to any literature-based review and cannot be avoided.

We did not investigate in detail how geographical and ecological overlap between host species affects pathogen sharing. We found that greater sympatry with other mammal species (defined as ≥20% area of target species range) was a positive predictor of viral sharing but was negatively correlated with bacterial sharing (Figure 4). Olival et al. (2017) concluded ≥20% was the minimum threshold for viral sharing, but it may not be appropriate for bacteria. Future work using generalized additive mixed models (GAMMs) will be necessary to properly control for autocorrelation in pathogen sharing networks (Albery et al., 2019), but we provide some thoughts here. Geographical overlap provides the necessary contact for host switching to occur (Davies & Pedersen, 2008), and some authors have claimed that the rate and intensity of contact may be “even more critical” than host relatedness in determining switching (Parrish et al., 2008). In support of this, ‘spillovers’ over greater phylogenetic distances are more common where vertebrates are kept in close proximity in zoos or wildlife sanctuaries (Kreuder Johnson et al., 2015). Similarly, although multi-host parasites generally infect hosts that are closely related rather than hosts with similar habitat niches (Clark & Clegg, 2017), ecology and geography have been found to be key factors influencing patterns of parasite sharing in primates (Cooper et al., 2012). Whereas, contact between two host species clearly provides a necessary but not sufficient condition for direct host switching, phylogenetic relatedness dictates the likely success of such a switch. Therefore, although the relative importance of phylogeny and geography may depend on the specific context, our observation of the strong dependence of pathogen sharing on phylogenetic distance across all vertebrates emphasises that this is the primary underlying biological constraint.

We have substantially improved on previous efforts to assess pathogen host range by using quantitative values based on alignment of multiple mitochondrial genes. However, our definition of species for pathogens remains somewhat arbitrary as it follows existing conventions. For example, in the *Mycobacterium tuberculosis* complex, the very closely related lineages *M. tuberculosis* (*n*=26 host associations), *M. bovis* (*n*=78), and *M. africanum* (*n*=3) are all treated as separate pathogens. Contrastingly, the extremely genetically diverse complex grouped under *Salmonella enterica* subsp. *enterica* (*n*=44 associations) is treated as a single pathogen. Exploring different measures of ‘phylogenetic scale’ to investigate these questions could help: phylogenetic scale is precisely defined for nested clades, but for non-nested and distantly-related clades it is less clear which measures to use (Graham, Storch, & Machac, 2018). Developing a parallel phylogenetic framework for pathogens to complement our host phylogenetic framework may be desirable, but challenging. An alignment of marker genes is tractable for bacteria (e.g. by using ribosomal proteins (Hug et al., 2016)), but more problematic for viruses, which have likely evolved on multiple independent occasions (Krupovic & Koonin, 2017). Tracing the ancestors of viruses among modern cellular organisms could represent another route to see if their host distribution reflects their evolutionary past. Potentially, an alignment-free genetic distance method could be used instead; as thousands more genomes become available for both pathogens and their hosts, such a method may be the optimal way to incorporate all known genomic information at a broad scale.

In conclusion, we have compiled the largest dataset of bacterial and viral pathogens of vertebrate host species to date. This is an important resource that has allowed us to explore different factors affecting the distribution of host range of vertebrate pathogens. While we are still some way off having a clear overall understanding of the factors affecting pathogen-host interactions, our results represent a substantial step in that direction. Maintaining such comprehensive datasets into the future is challenging but important, in order to ensure that all available knowledge is synthesized — rather than drawing conclusions only from well-studied pathogens, which likely represent the exceptions and not the norm.

## Acknowledgements

We acknowledge financial support from the European Research Council (ERC) (grant ERC260801—BIG_IDEA to FB). DD and CD acknowledge the support of the Swiss National Science Foundation (grant 150654). We would also like to acknowledge the many public databases used in the construction of our own database and thank all creators. We thank Olival and colleagues for releasing comprehensive code for their reproducible analyses, allowing us to adapt it for our own purposes and easily compare results.

## Data Accessibility Statement

Associated data and analysis code is available as a Github repository (https://github.com/liampshaw/Pathogen-host-range) and also archived on figshare (doi: <u>10.6084/m9.figshare.8262779</u>). This includes: the pathogen-host association database; the host phylogenetic tree; other datasets derived from them; and an Rmarkdown notebook which reproduces all analyses in this paper.

## Image credits

**Figure 1**. Icons were downloaded from FlatIcon (flaticon.com) and were made by Maxim Basinski (tick/cross symbols) and Chanut (document symbol). Pathogen images (influenza, bacterium, adenovirus, HIV) were from the Bacteriology Virology image set from Servier Medical Art (smart.servier.com). Silhouettes were downloaded from PhyloPic (phylopic.org) and were made available under the Public Domain Dedication license (human and monkey) or a Creative Commons Attribution license: falcon (by Liftarn, CC BY-SA 3.0: <u>creativecommons.org/licenses/by-sa/3.0</u>), opossum (by S. Werning, CC BY 3.0: <u>creativecommons.org/licenses/by/3.0</u>).

**Figure 3**. Silhouettes were downloaded from PhyloPic. Images were either made available under the Public Domain Dedication license (Chiroptera, Cingulata, Peramelemorphia, Primates, Scandentia) or a Creative Commons Attribution license: Carnivora and Rodentia (by R. Groom, CC BY 3.0 and CC BY-SA 3.0 respectively); Didelphimorphia, Pilosa, Lagomorpha, and Diprotodonia (by S. Werning, CC BY 3.0); Euliptyphla (by C. Rebler, CC BY-SA 3.0); Perissodactyla (by J. A. Venter, H. H. T. Prins, D. A. Balfour & R. Slotow and vectorized by T. M. Keesey, CC BY 3.0); Probscidea (by T. M. Keesey, CC BY 3.0); Cetartiodactyla (by Zimices, CC BY 3.0).

**Figure 6.** Silhouettes were downloaded from PhyloPic. Images were either made available under the Public Domain Dedication license (Chiroptera) or the Creative Commons Attribution license: Primates (by B. Bocianowski, CC BY 3.0), Didelphimorphia (by S. Werning, CC BY 3.0), Falconiformes (by Liftarn, CC BY 3.0).

## Author contributions

Using the CRrediT taxonomy, the contributions of all authors to different stages of the project were as follows: conceptualization (FB, ADW, LPS), data curation (ADW, DD, MM, GP, LPS), formal analysis (ADW, LPS, DD), funding acquisition (FB, CD), investigation (ADW, LPS, DD), methodology (ADW, LPS, DD, CD, FB), software (ADW, LPS, DD), supervision (CD, FB), visualization (ADW, DD, LPS), writing – original draft (LPS, ADW, FB), writing – review & editing (LPS, ADW, DD, CD, FB).

