## Supplementary Material for "The phylogenetic range of bacterial and viral pathogens of vertebrates"

**Supplementary Code Repository:** available at <https://github.com/liampshaw/Pathogen-host-range> (archived version at: <https://doi.org/10.6084/m9.figshare.8262779>). Contains an Rmarkdown notebook which collates code to produce figures and tables.

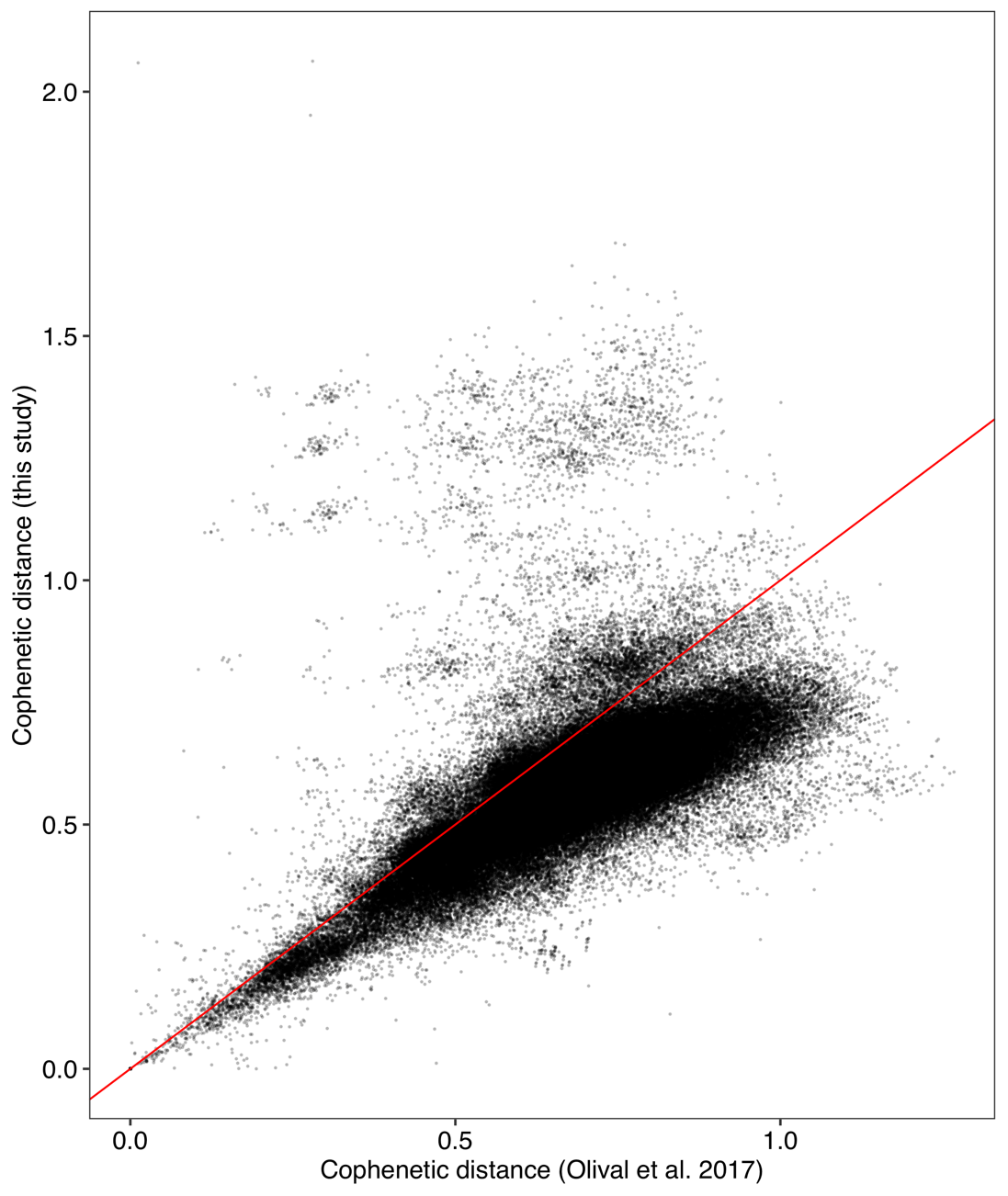
**Supplementary Figure 1. Correlation of cophenetic distances between this study (y-axis) and Olival et al. (2017).** Cophenetic distance shown for all pairwise comparisons of vertebrate host species shared between datasets (*n*=551) for the *cytb* tree published by Olival et al. and the multi-mitochondrial gene tree used in this study.

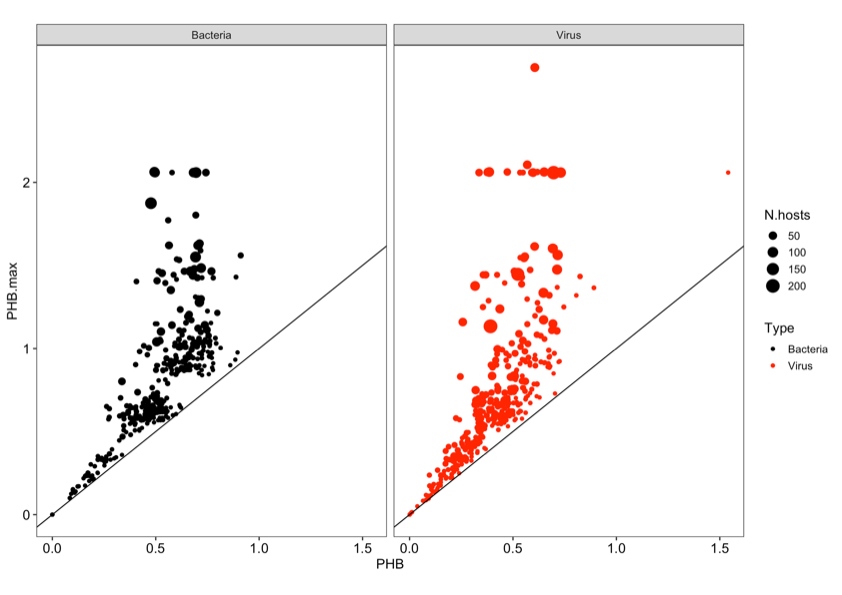
**Supplementary Figure 2. Correlation of mean and maximum PHB for pathogens.** Correlation is shown for bacteria (left) and viruses (right).

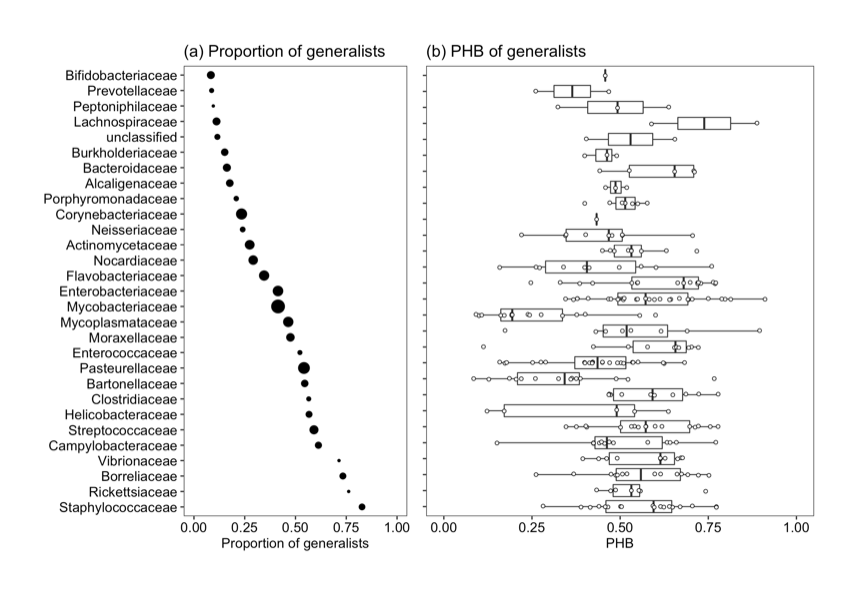
**Supplementary Figure 3. (a) Proportion of generalists and (b) PHB of generalist pathogens by bacterial family.** Only bacterial families with >20 pathogen species in the association database are shown. Families are ordered by the proportion of generalists. There is no clear association between the proportion of generalists within a family and how wide-ranging those generalists are.

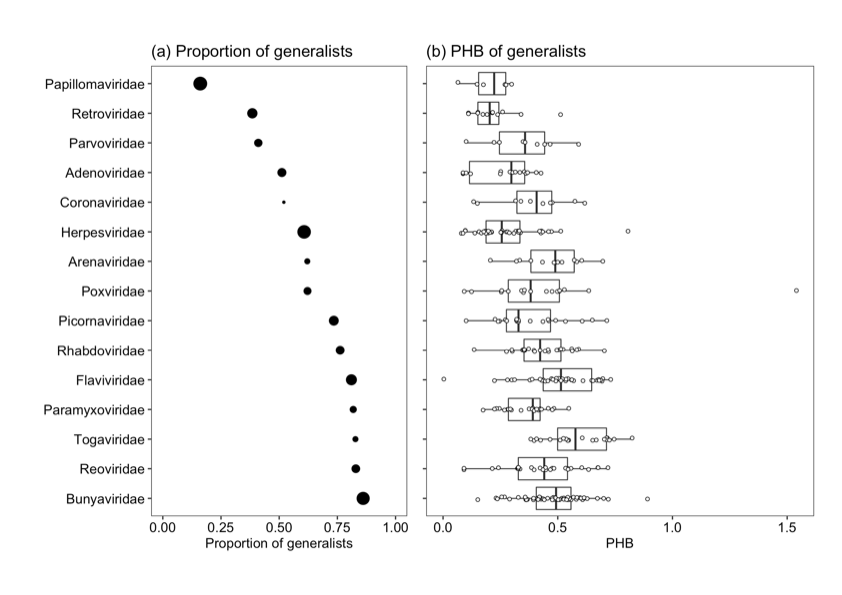
**Supplementary Figure 4. (a) Proportion of generalists and (b) PHB of generalist pathogens by viral family.** Only viral families with >20 pathogen species in the association database are shown. Families are ordered by the proportion of generalists.

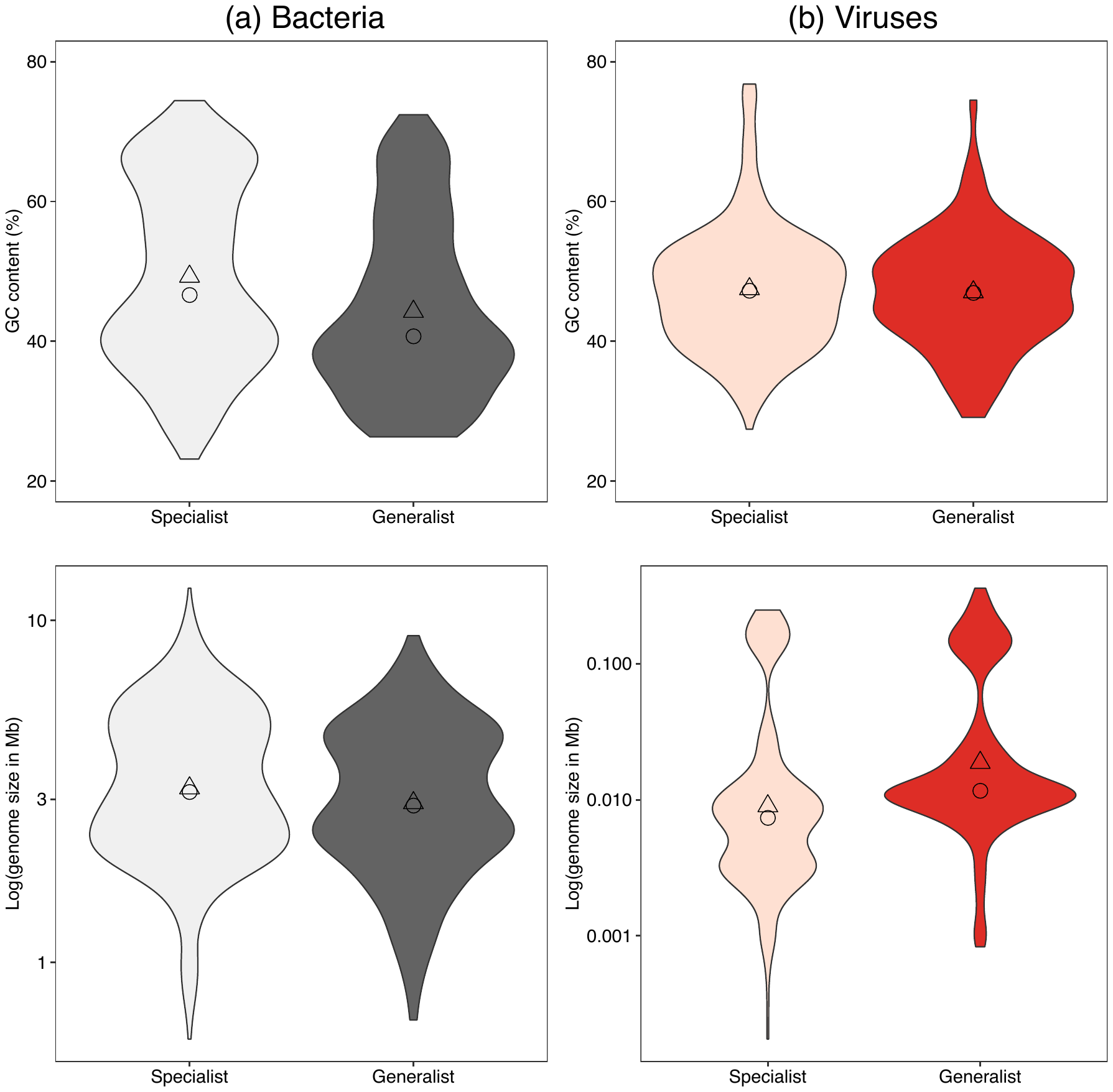

**Supplementary Figure 5. Pathogen genome GC content and size for specialist and generalist pathogens.** Note the log-scale for the y-axis in the lower half of the figure. Distributions are shown for specialists (light colours) and generalists (dark colours) with their median (circle) and mean (triangle).

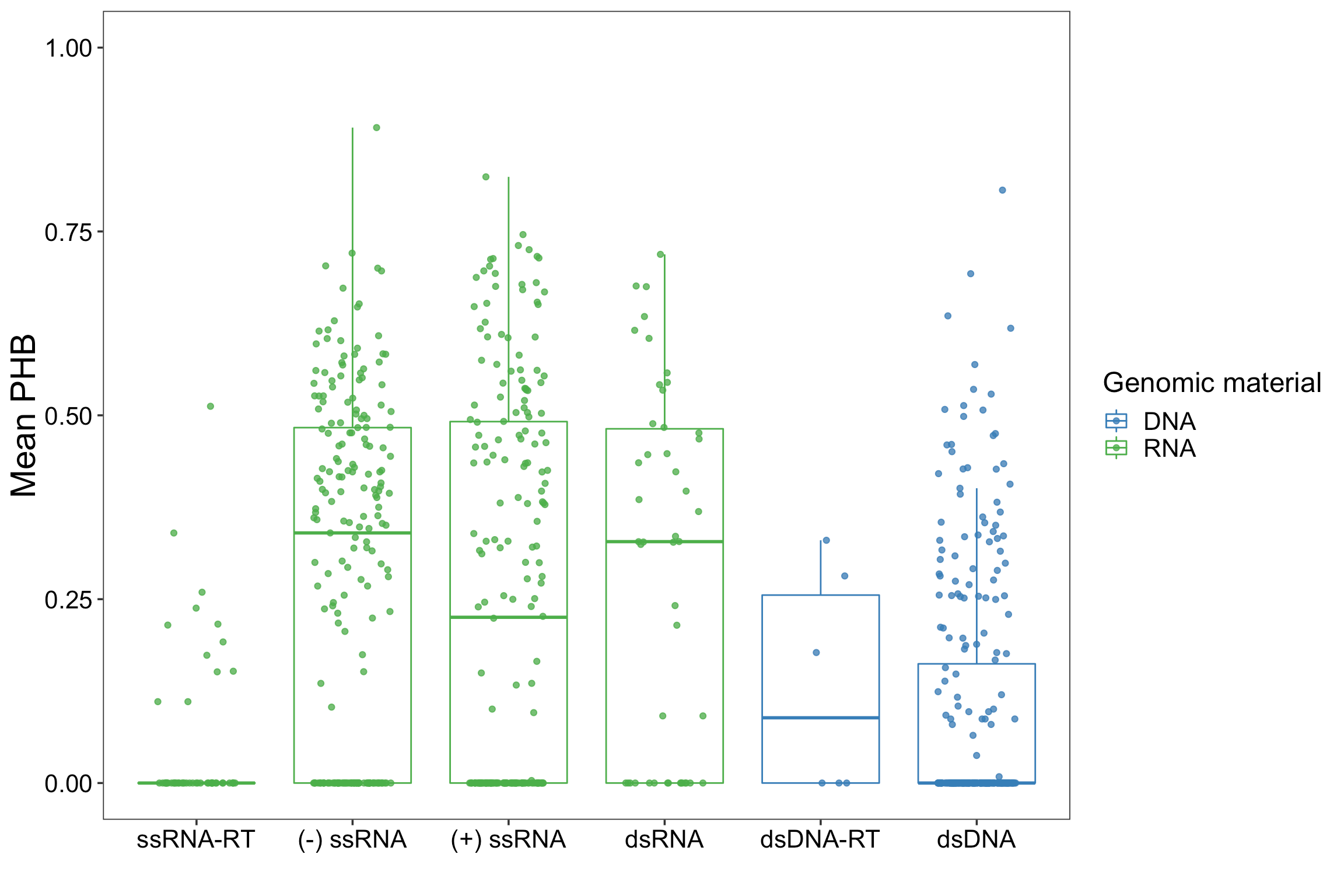
**Supplementary Figure 6. Viral genome type is associated with host range.** RNA viruses have a greater median mean PHB than DNA viruses. Subgroups shown are the Baltimore classification.

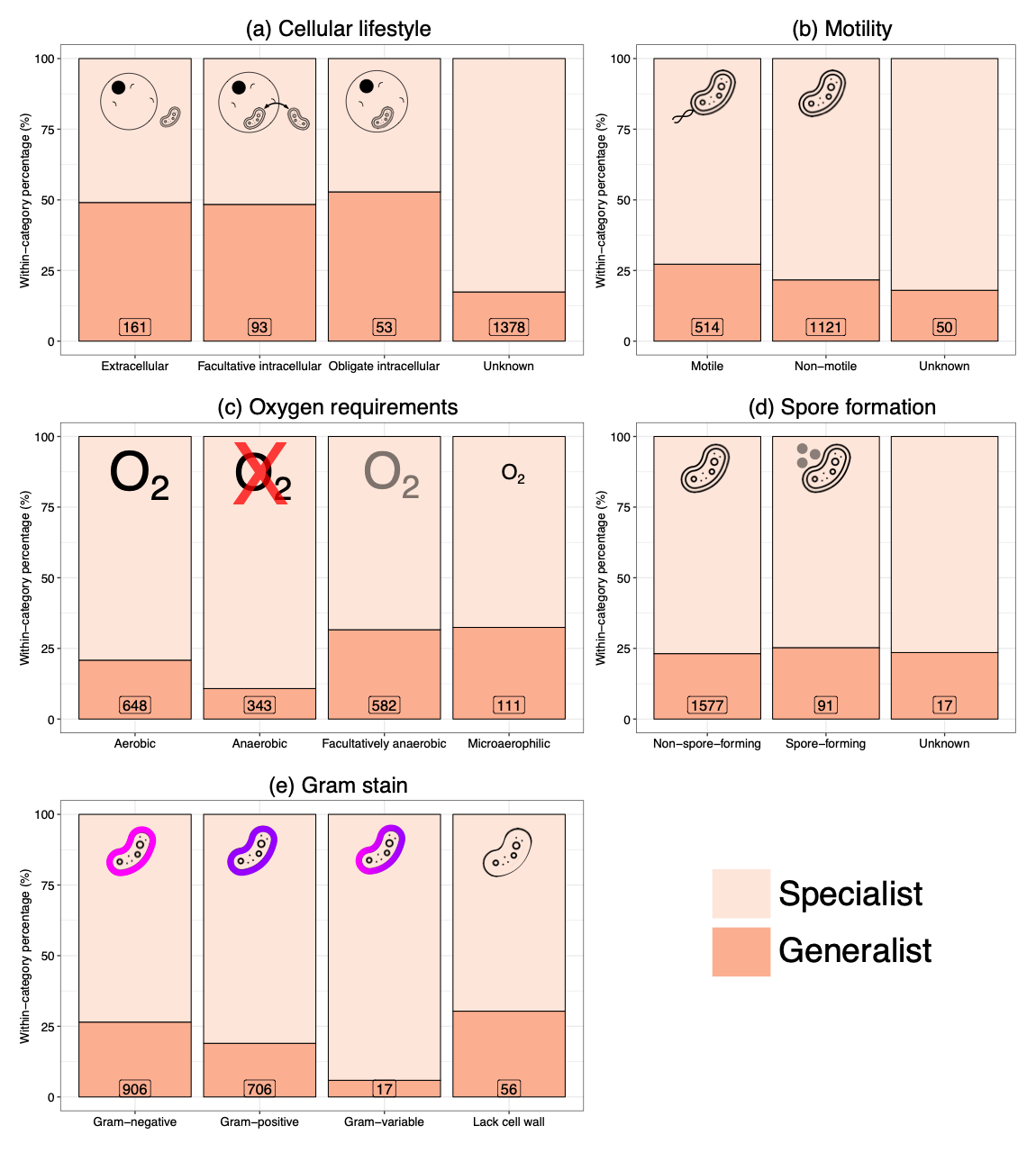

**Supplementary Figure 7. Bacterial lifestyle factors and pathogen range.** Proportion of specialists (light pink) and generalists (dark pink, PHB>0) for different categories of bacterial lifestyle: (a) cellular lifestyle, (b) motility, (c) oxygen requirements, (d) spore formation, and (e) Gram stain. ‘Unknown’ can also mean ‘not applicable’.

**Supplementary Table 1. Vector-borne pathogens are more likely to be generalists.** Number of bacteria and viruses with known invertebrate vectors.

| Viruses | Specialist | Generalist |
| --- | --- | --- |
| Not vector-borne | 469 | 271 |
| Vector-borne | 50 | 117 |
| Bacteria | | |
| Not vector-borne | 1240 | 337 |
| Vector-borne | 53 | 52 |

**Supplementary Table 2. Viruses with an RNA genome and larger genome size have a greater host range.** Having an RNA genome and a larger genome were both significantly associated with greater mean PHB (*p*<0.001 for both variables) with a non-significant interaction between them (*p*=0.36). Interestingly, genome size was not significantly associated with greater PHB in a univariate model (Supplementary Text 1).

|  | Coefficient (s.e.) | p |
| --- | --- | --- |
| Intercept | 0.048 (0.015) | 0.002 |
| RNA genome | 0.188 (0.031) | <0.001 |
| Genome size | 0.661 (0.148) | <0.001 |
| Interaction | 1.742 (1.919) | 0.365 |

**Supplementary Table 3. Bacterial motility and cellular lifestyle are not associated with greater host range.** Combining motility and cellular proliferation in a linear model suggests that neither variable is associated with greater mean PHB. Univariate linear models and a linear model with an interaction term (Supplementary Text 1) give the same conclusion.

|  |  | Coefficient (s.e.) | p |
| --- | --- | --- | --- |
| Intercept | | 0.264 (0.028) | <0.001 |
| Cellular lifestyle | Facultative intracellular | -0.042 (0.038) | 0.278 |
|  | Obligate  intracellular | -0.004 (0.051) | 0.933 |
| Motility | Not-motile | 0.025 (0.037) | 0.502 |
